# The role of actin and myosin II in the cell cortex of adhered and suspended cells

**DOI:** 10.1101/2021.08.03.454901

**Authors:** D.A.D. Flormann, K.H. Kaub, D. Vesperini, M. Schu, C. Anton, M.O. Pohland, L. Kainka, G. Montalvo Bereau, A. Janshoff, R.J. Hawkins, E. Terriac, F. Lautenschläger

## Abstract

Adhesion induces dramatic morphological and mechanical changes to cells, which are reflected by changes to the actin cortex. Among the many different proteins involved in this sub-membranous layer, motor proteins (e.g., nonmuscle myosin II [NMII]) and actin nucleators (e.g., Arp2/3, formins) are known to have significant influences on its dynamics and structure. The different roles of NMII, Arp2/3, and formins in the dynamics, structure, and mechanics of the actin cortex depend on the adhesion state of the cell. In this study, we unravel the interplay between the dynamics, structure, and mechanics of the actin cortex in adhered cells and in cells in suspension. We show that treatments with extrinsic cellular perturbants lead to alterations of all three properties that are correlated. However, intrinsic actin cortex variations between different cell adhesion states lead to unexpected correlations. Surprisingly, we find that NMII minifilaments have a minor influence on the actin cortex. Using new microscopy techniques, we show that NMII minifilaments are not localized within the actin cortex, as previously thought, but concentrated in a layer beneath it. Our treatments affecting Arp2/3 and formin reveal correlations between the actin cortex characteristics. Our data build towards a comprehensive understanding of the actin cortex. This understanding allows the prediction and control of cortical changes, which is essential for the study of general cellular processes, such as cell migration, metastasis, and differentiation.

## Introduction

Actin is the most abundant protein in eukaryotic cells [1], and its combination with microtubules and intermediate filaments defines the cytoskeleton. The main structure responsible for the mechanical properties of cells is the actin cortex, which is assembled directly under the plasma membrane. As the actin cortex is such a pivotal cellular element, it has stimulated a lot of studies, especially for its roles in cell mitosis, migration, and differentiation [2–4]. Moreover, a direct relation between actin concentration and actin network stiffness has been shown [5].

The structure of the actin cortex remains poorly understood from both the molecular and architectural points of view. Along with the main actin component, there are many other molecules that contribute to the properties of the actin cortex. The molecular motors such as nonmuscle myosin II (NMII) provide contractile forces, while a combination of actin nucleators, (de-)stabilizers, and capping proteins controls the rate of actin polymerization. This combined regulation allows the structure of the actin cortex to adapt to both intrinsic (e.g., cell rounding at entry into mitosis) and extrinsic (e.g., osmotic shock) changes.

One particular change that some cell types can experience in vivo involves their adhesion state. Metastatic cells, for example, detach from multicellular aggregates to move through the body, either individually or collectively with other tumor cells. These changes in their adhesion state involve alterations to the actin cortex. This has been shown by a reported decrease in the stiffness of cancer cells, compared to their noncancer originals [6, 7].

The actin cortex is influenced by a broad range of proteins, and here we focus in particular on NMII, Rho-associated protein kinase (ROCK) and the actin nucleators Arp2/3 and formin. NMII is considered to be the main molecular motor that contributes to cellular mechanics. However, the influence of NMII in both adhered and suspended cells mechanical properties remains controversial in the literature [3, 4, 8–12]. ROCK is a known activator of NMII but also influences a broad range of proteins and is involved in several cellular processes, such as apoptosis and cell migration. As ROCK is down-regulated in suspended cells, it has a crucial role in cell adhesion state transitions, such as during metastasis [13].

Arp2/3 is a seven-protein complex that promotes nucleation of new actin filaments as side (daughter) filaments of the primary (mother) filaments, which results in branched actin networks [14]. On the other hand, formin binds to the barbed ends of established actin filaments, where it accelerates actin polymerization, and hence it is an important part of the machinery that rearranges the actin cortex in cells [15]. It has been suggested that mDia is the main active formin in the actin cortex [16]. Arp2/3-mediated and formin-mediated nucleation and polymerization are not mutually exclusive, and the balance in the recruitment of each of these nucleation factors is orchestrated through different nucleation-promoting factors. Indeed, the level of activity of each of these nucleation factors is a strong determinant of the structure of the actin cortex.

Here, we propose a novel paradigm for the mechanics of cell cortices with far-reaching implication of a comprehensive understanding how cells respond and adapt to external cues (challenges) and generate complex function by driving an active matter network far from equilibrium.

## Results and discussions

Immortalized RPE1 cells are of epithelium origin, although they also show mesenchymal characteristics, such as high levels of N-cadherin and low levels of E-cadherin [17]. Consequently, these cells share similarities with metastatic cancer cells that transition between the epithelial and migratory phenotypes during the initial metastasis.

To better understand the dynamics, structure, and mechanics of the actin cortex, we focused on the most ‘extreme’ of the cell adhesion states: fully adhered cells (i.e adhered for more than 12 hours), and cells in suspension (Figure 1A, B). For adhered cells, it is known that their mechanical responses differ across cell regions [8, 18], and therefore we considered the adhered cell state in terms of two regions: the actin cortex over the nucleus and in the perinucleus. We define the perinucleus as between the nuclear edge and the cell edge (compare Figure S9 and S10) [19]. In contrast, the actin cortex of spherical, suspended cells was investigated without any regional separation (Figure S10). We define these measures of the dorsal actin cortex over the nucleus, in the perinucleus, and in suspended cells as the NPS (‘Nucleus’, ‘Perinucleus’, ‘Suspended [cells]’) states.

**Figure 1:**
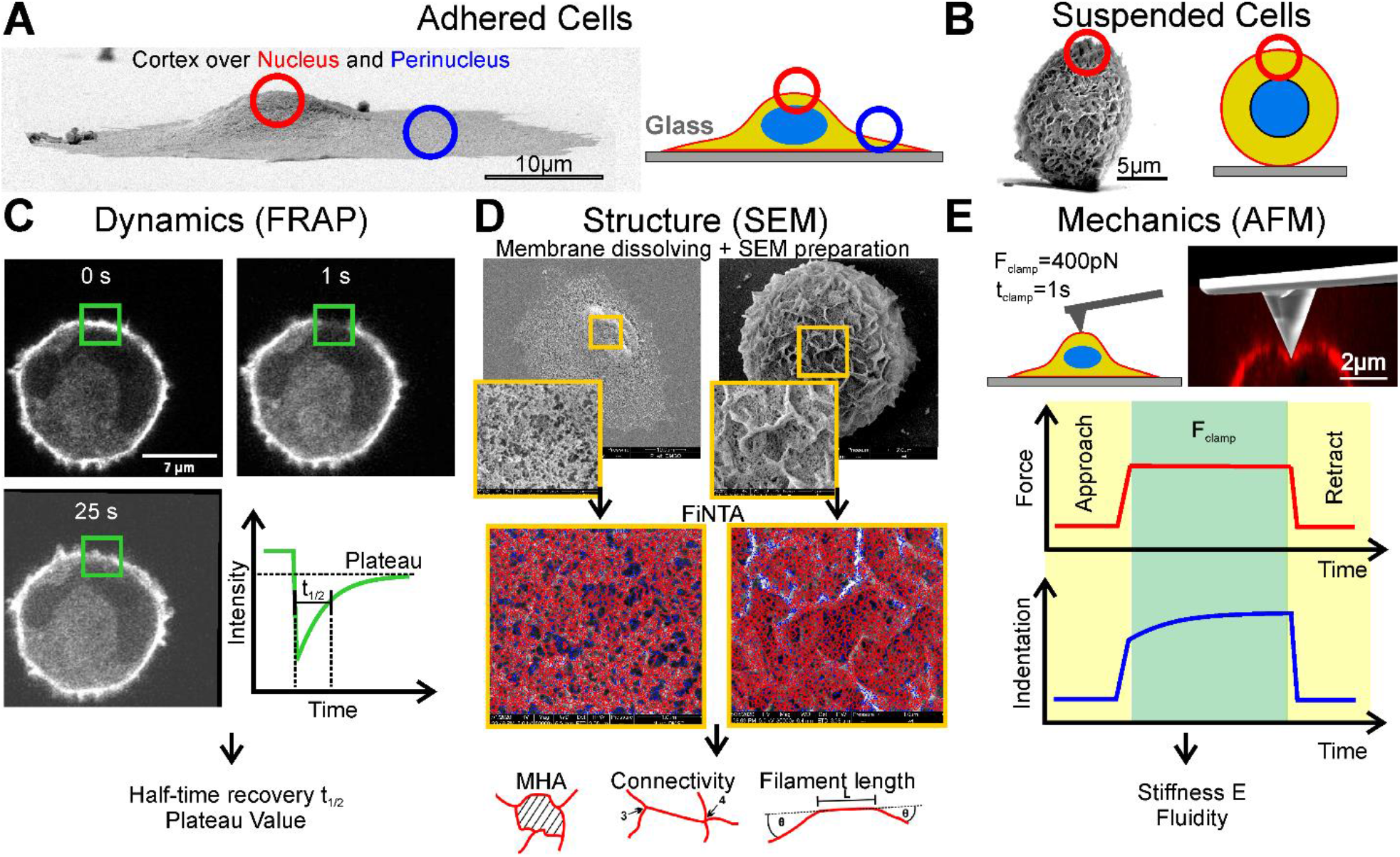
Overview of cell states and investigation methods. The actin cortex of adhered hTERT-RPE1 cells was measured over the nucleus and the perinucleus illustrated with SEM side-view and scheme (**A**). The actin cortex of suspended was measured at the top (SEM and AFM) or at the equatorial plane (FRAP) (**B**). FRAP investigation were performed using a LSM in AiryScan-mode (**C**). SEM preparation and exemplary actin fiber tracing using FiNTA (**D**). AFM investigations were performed using creep-compliance measurement with a force clamp of 400pN over 1s. Photomontage of fluorescence (cell, actin in red) and SEM imaging (cantilever) illustrates the cortex deformation (confocal z-reconstruction while AFM was mounted on the confocal microscope) upon indenting a soft suspended cell using AFM, while cantilever was separately images with SEM (**E**).

Three particular aspects were investigated here: the actin dynamics, the actin cortex structure, and the cellular mechanics, which we refer to as the DSM (‘Dynamics’, ‘Structure’, ‘Mechanics’) measures. These were quantified through the following techniques: fluorescent recovery after photobleaching (FRAP) for the dynamics of the actin cortex within the cells, as the actin half-time recovery (t_1/2_) and intensity plateau value (Figure 1C); scanning electron microscopy (SEM) for the structure of the cellular actin cortex, to measure the mesh hole area (MHA), connectivity, and filament length (Figure 1D); and local creep compliance atomic force microscopy (AFM) to measure the cellular mechanics as cell stiffness (E) and cell fluidity (Figure 1E). The main focus for these DSM measures were t_1/2_ (for actin dynamics), MHA (for actin structure), and E (for actin mechanics).

These approaches led to the ‘basic dataset’ that is referred to throughout this study (Figure 2). In the next three sections, the basic dataset was used to investigate extrinsic and intrinsic correlations between the DSM measures per se and the NPS states per se. Then in the last two sections, the influence of various cytoskeletal perturbants was investigated, in terms of their effects on the DSM measures for the different NPS states, the localization of NMII, Arp2/3, and formin.

**Figure2:**
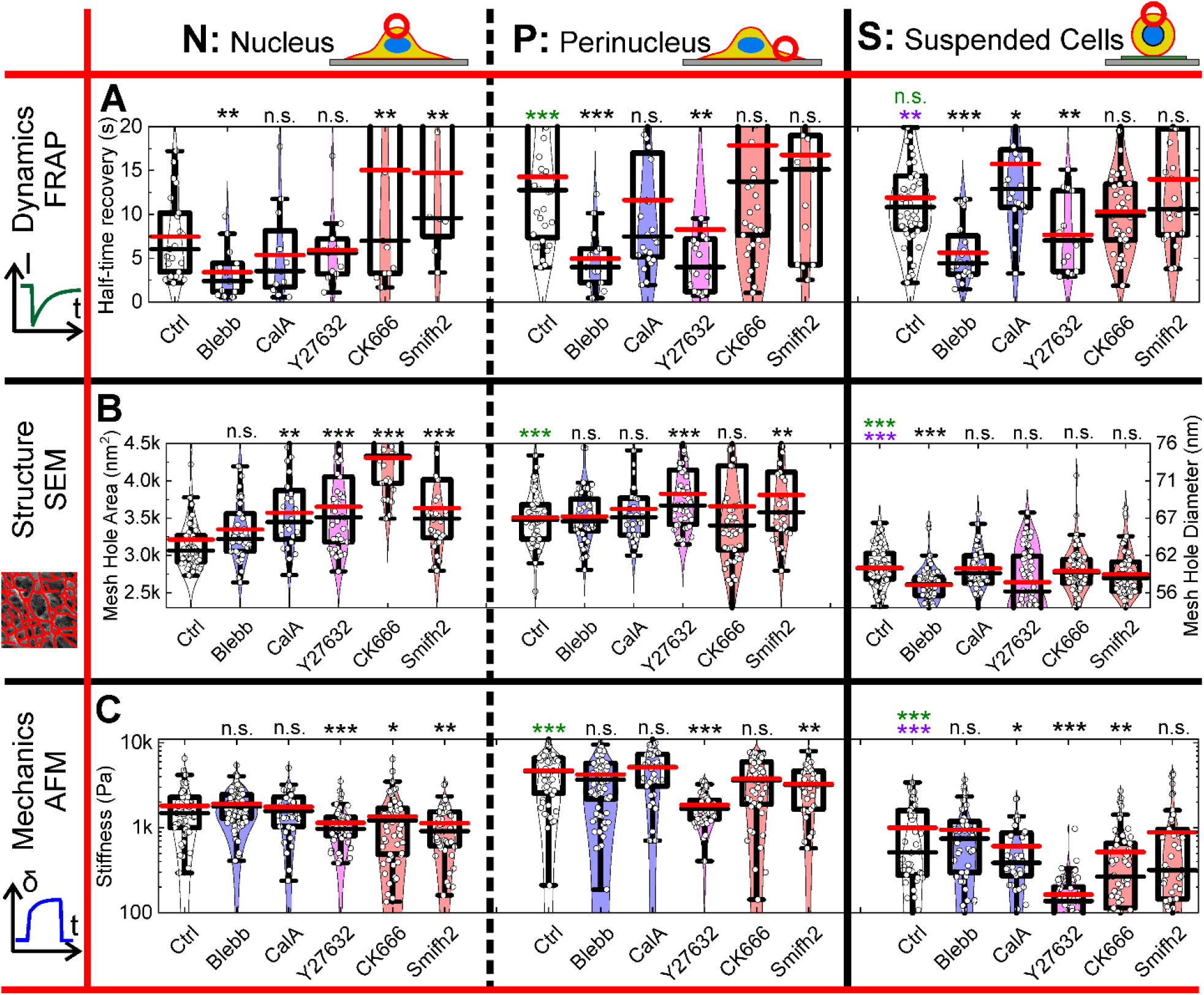
Structure, mechanics and dynamics of the cellular cortex of adhered (nucleus and perinucleus) and suspended hTERT-RPE1 cells. Half-time recovery t_1/2_ was extracted from Fluorescence Recovery After Photobleaching measurements with a Laser Scanning confocal Microscope LSM (**A**). The MHA of the actin cortex was quantitatively analyzed using Scanning Electron Microscopy images (**B**). The stiffness was quantitatively analyzed employing creep compliance measurements using Atomic Force Microscopy (**C**). Median values are marked in red, mean values in black. Star method is representing statistical Welch-corrected t-tests. Black stars compare pharmagolocial perturbants with the Ctrl for each panel. Green stars compare controls to Nucleus Ctrl, purple stars compare Suspended Ctrl to Perinucleus Ctrl. n.s.: not significant, *: p<0.05, **: p<0.01, ***: p<0.001. Cell numbers n are in the order Ctrl, Blebb, CalA, Y27632, CK666, Smifh2. *Dynamics (FRAP)*: Nucleus: n= 21, 12, 12, 11, 13, 8; Perinucleus: n= 32, 13, 24, 21, 42, 13; Suspended cells: n= 50, 20, 20, 14, 40, 23; *Structure (SEM)*: Nucleus:n= 63, 39, 24, 41, 26, 31; Perinucleus: n= 57, 40, 30, 46, 54, 33; Suspended cells: n= 66, 75, 44, 84, 70, 70; *Mechanics (AFM)*: Nucleus: n= 53, 88, 38, 51, 63, 45; Perinucleus: n= 52, 73, 34, 49, 47, 31; Suspended cells: n= 42, 47, 35, 49, 57, 41.

### DSM measure correlations: Extrinsic alterations to the actin cortex unravel clear correlations between actin dynamics, structure, and mechanics

Considering the individual DSM measures (i.e., t_1/2_, MHA, E), the basic dataset indicated separation between the cell adhesion (NPS) states (Figure S1A-C). As ROCK is naturally down-regulated in suspended cells and has severe downstream effects on both NMII and actin, correlations between the DSM measures were calculated without and with Y-27632, a ROCK inhibitor (Figure 3, black, magenta, respectively) [13, 20]. Pearson correlation coefficients were used as the measure of correlation.

**Figure 3:**
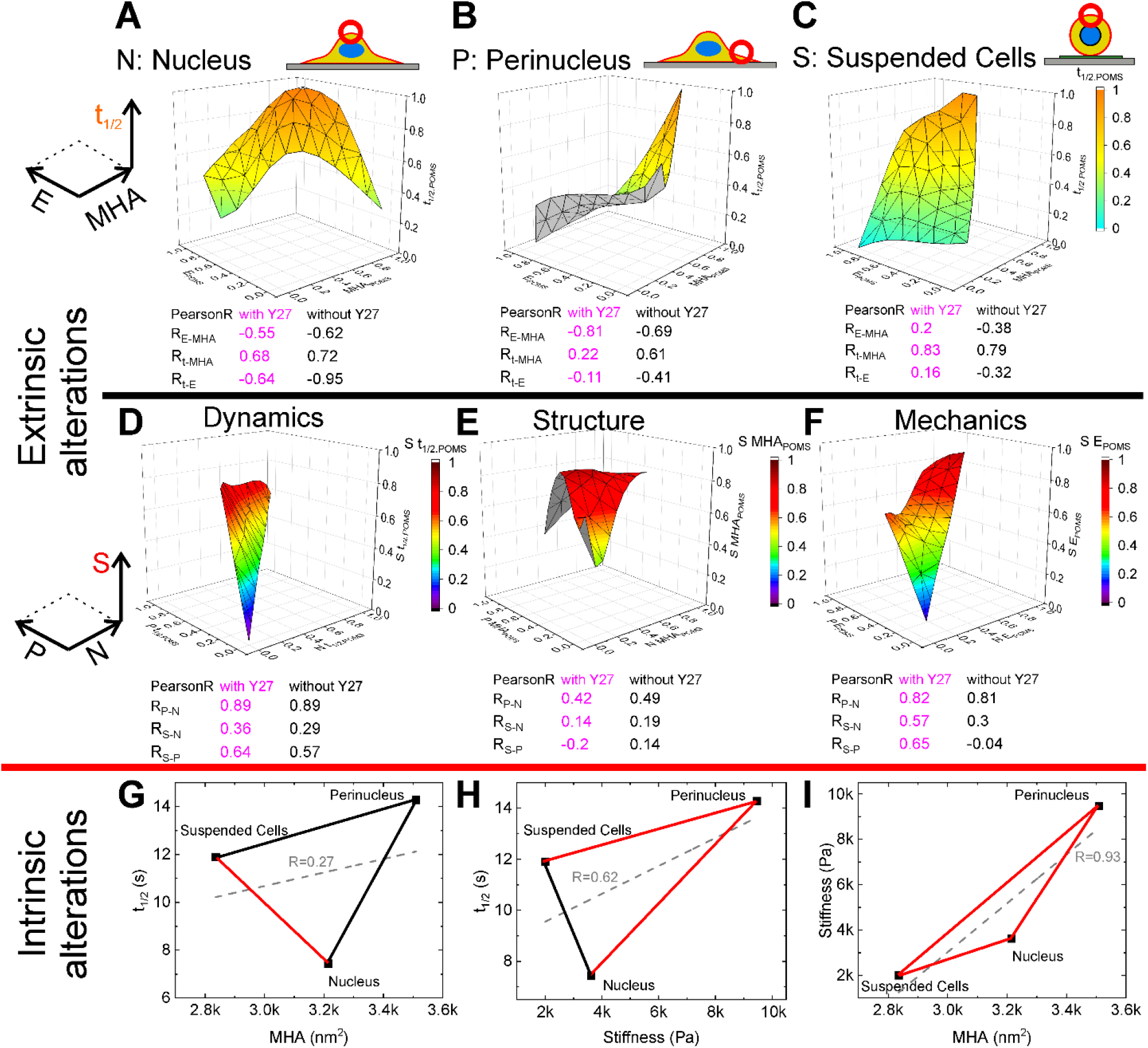
Extrinsic and intrinsic correlations. 3D correlation diagrams between dynamics, structure and mechanics upon extrinsic cortex alterations for nucleus (**A**), perinucleus (**B**) and suspended cells (**C**). 3D correlation diagrams upon extrinsic cortex alterations between nucleus, perinucleus and suspended cells for dynamics (**D**), structure (**E**) and mechanics (**F**). 2D planes of each 3D diagram are depicted in Figures S2 (for A-C) and S3 (for D-F). Surfaces in 3D diagrams are the result of extrapolation between data points (see Figure S2 and S3). Pearson R correlations coefficients are calculated with (magenta) and without (black) Y-27632 taken into account (**A-F**). 2D “correlation” diagrams upon intrinsic cortex alterations for Ctrl cells (compare Figure 2) (**G-I**). Red connections represent opposite algebraic signs of Pearson R’s compared to A-C. Black connections represent identical algebraic signs of Pearson R’s compared to A-C. For clarity error bars are not shown here, since they are depicted in Figure 2. Gray dashed lines represent linear approximations of all three cell states/regions with Pearson R correlation coefficient. For all diagrams data from Figure 2 were normalized using POMS normalizations (see Figure S1). Each 3D diagram consists of 18 data points (means of Figure 2) with Y27632 and 15 without Y27632. The total number of cells n used to achieve these were: (A) n= 639; (B) n= 691; (C) n= 847; (D) n= 389; (E) n= 893; (F) n= 895 with Y27632 and (A) n= 536; (B) n= 575; (C) n= 700; (D) n= 343; (E) n= 722; (F) n= 740 without Y27632. (G) Nucleus: n= 84; Perinucleus: n= 89; Suspended cells: n= 116; (H) Nucleus: n= 74; Perinucleus: n= 84; Suspended cells: n= 92; (I) Nucleus: n= 116; Perinucleus: n= 109; Suspended cells: n= 108.

First, we investigated each NPS state separately. This provided one parameter space diagram for each NPS state individually, for comparisons of the focus parameters t_1/2_, MHA, and E (Figure 3A-C). For clear presentation, we depict all of the planes of the three-dimensional diagrams separately in two dimensions (Figure S2B-D).

This showed that the correlations between DSM measurements in adhered cells (both over the nucleus and in the perinucleus) were independent of ROCK inhibition, with t_1/2_ positively correlated with MHA and negatively correlated with E (Figure 3A, B). Also, E was negatively correlated with MHA, as has been predicted in vitro [5, 21]. For suspended cells, these correlations were similar in the re-normalization without ROCK inhibition (Figure 3C). In contrast, with ROCK inhibition, only t_1/2_ and MHA remained strongly correlated, with no correlations between any of the other parameters (−0.2 ≤ R ≤ 0.2). We postulate that this is due to down-regulation of ROCK in suspended cells [13]. In absence of an active substrate, Y27632 affinity might be focused on other substrate such as PKCe [22] which will interfere with other cellular pathways than those in focus in this study.

In summary, inhibition of ROCK decoupled the correlations involving cell stiffness only for suspended cells, which will be discussed further later. We hypothesize that the formation of the actin cortex begins with its dynamics, which influences its structure. This has direct consequences on cell stiffness. For the DSM measures, this means that an increase in t_1/2_ (i.e., slower actin polymerization) will increase MHA, which in turn reduces E. This appears reasonable, as an infinite t_1/2_ would lead to complete depolymerization of the actin cortex (i.e., infinite MHA). This infinite MHA would lead to significantly lower E, as observed by treatments with the actin polymerization inhibitor latrunculin A, and by the combination of the actin polymerization inhibitor CK666 and the actin nucleation inhibitor Smifh2 (Figure S7). The effects of the combination of these last two perturbants were excluded from the main analysis here, due to the severe cell shrinkage that they caused (compare Figure S9A, B).

### NPS state correlations: Extrinsic alterations to actin DSM measures in one adhesion state correlate only partly with other adhesion states

After establishing the correlations between DSM parameters, we asked if one parameter, e.g. t_1/2_, is extrinsically altered in adhered cells, does the same parameter (here t_1/2_) alter in the same manner upon identical extrinsic alterations in suspended cells? This question was asked for each of the DSM measures.

As shown in Figure 3D, E, perturbations of the cortex in adhered cells resulting in alteration to any of the DSM measures over the nucleus always correlated positively with the corresponding DSM measures of the perinucleus (Figure 3D-F; R_P-N_). Further, alterations to t_1/2_ and E over the nucleus correlated with similar alterations of these two parameters within the actin cortex of suspended cells (Figure 3D, F; R_S-N_). However, no such correlation was observed for the structure of the actin cortex (Figure 3E; R_S-N_). Comparing suspended cells with the perinucleus of adhered cells, t_1/2_ behaved similarly for both (Figure 3D; R_S-P_), while MHA behaved fundamentally differently (Figure 3E; R_S-P_). Surprisingly, the mechanics were only correlated between the perinucleus and suspended cells if Y-27632 was included (i.e., ROCK was inhibited), with no correlation seen without Y-27632 (Figure 3F; R_S-P_).

For simplification, we also summarized the DSM measures for the nucleus and perinucleus as the adhered cells collectively, as “NP”, and compared this directly with suspended cells (Figure S3M-O). As expected, the actin dynamics of adhered cells correlated with the actin dynamics of suspended cells, while the actin structure was not correlated between these cell adhesion and suspension states. Including ROCK inhibition data using Y-27632 led to mechanical correlations between adhered and suspended cells, while this was not the case when those data were excluded from the correlation calculations. Consequently, t_1/2_ is independent of cell adhesion state, MHA depends strongly on cell adhesion state, and the cellular mechanics depend on ROCK.

As the DSM measures are conserved within the adhered cells (i.e., N versus P) and the correlations between adhered and suspended cells appear to depend partly on ROCK, we hypothesized that the uncorrelated results are due to the natural down-regulation of ROCK in suspended cells [13]. Further details are given in support of this in the Supplementary Information, along with the two-dimensional representations in Figure 3D-F.

### Intrinsic actin cortex alterations induced by cell adhesion state transition reveal the underlying biological processes

To investigate the intrinsic actin cortex alterations induced by the cell NPS ‘adhesion transition’, we analyzed the data for the untreated cells (Figure 2, Ctrl). In contrast to the DSM correlations in Figure 3A-C, these intrinsic alterations to the actin cortex primarily led to unexpected correlations (Figure 3G-I).

First, smaller MHA was not correlated with higher E (Figure 3I), as was seen for the extrinsic alterations (Figure 3A-C), but the opposite i.e. larger MHA correlates with higher E. This implies that cell stiffness E is influenced by different factors. Secondly, t_1/2_ and MHA were correlated, as expected, although not for the actin cortex of suspended cells compared to that over the nucleus of adhered cells (Figure 3G). Thirdly, t_1/2_ was correlated with stiffness E, as expected, only for the actin cortex of suspended cells compared to that over the nucleus only of adhered cells (Figure 3H).

These observations show the important role of the underlying biological processes upon transition between the NPS states. We hypothesized that these counter-intuitive effects are explained by the three following aspects (see Supplementary Information for further details):

1. The natural down-regulation of ROCK in suspended cells leads to increased cofilin activity, and therefore to weakening of the actin cortex [13, 23]. Although ROCK influences a broad range of proteins, we focused on its effects on cofilin (see Supplementary Information). We hypothesized that the actin cortex of suspended cells consists of shorter and more weakly bound actin filaments compared to the actin cortex of adhered cells. We believe that this specific architecture of the actin cortex of suspended cells directly influences the DSM measures i.e. we propose that the smaller MHA is due to shorter filaments and lower stiffness is due to filaments being less strongly connected.
2. High actin stress within the perinucleus due to stress fibers and actin bundles leads to increased cell stiffness and a ‘stretched’ actin network (i.e., larger MHA) compared to over the nucleus and to suspended cells. Although MHA was larger for the perinucleus, the stiffness was higher in this region than over the nucleus and for suspended cells. Intuitively, we would have expected that the smaller the MHA, the greater the stiffness. As we observed the opposite, we hypothesize that the increase in the stiffness for the perinuclear region compared to over the nucleus and to suspended cells is mainly related to the presence of stress fibers and actin bundles, which are known to be under high stress (Figure S9E). This argument extends to the comparison between over the nucleus and in suspended cells, as the actin cortex over the nucleus is connected to the actin cortex of the high-stress perinucleus, whereas there is no evidence for a similar high stress reqion in suspended cells, for which we observed the smallest MHA and the lowest stiffness (compared to over the nucleus and to the perinucleus).
3. The membrane curvature increases from the perinucleus, to over the nucleus, to suspended cells. The membrane curvature influences actin polymerization via several pathways (e.g., increased N-WASP and Arp2/3 recruitment with increased membrane curvature) [24, 25]. As all of the NPS states have different membrane curvatures, we hypothesized that these curvature changes lead to the different correlations of the intrinsic to extrinsic cortical alterations.

As both the underlying biological processes and the extrinsic actin cortex alterations will also rely on important key proteins of the actin cortex within each NPS state, we next investigated the physical locations of NMII, Arp2/3, and formin.

### Nonmuscle myosin II minifilaments are positioned beneath the actin cortex

Cytoskeleton perturbants that target NMII, such as blebbistatin and calyculin A, had little effect on the structure and stiffness of the actin cortex, but had large effects on the actin dynamics (Figure 2). This was surprising, since we had expected NMII minifilaments to have a stronger impact on the actin cortex mechanics. We therefore questioned whether NMII minifilaments are localized in the cortex. To test this, we performed high resolution expansion microscopy in combination with Zeiss Airy scans for the NMII localization close to the cortex. This has a, theoretical resolution of ~22 nm, which is 4.5-fold that for conventional confocal microscopy [26].

To acquire high-resolution images of the ventral and dorsal sides of the actin cortex of these cells, they were first embedded in hydro-gel, and then slices of the gel (containing the cells) were flipped directly onto a glass dish (Figure 4A, B). This enabled the acquisition of high-resolution expansion microscopy images of the ventral and dorsal sides of the actin cortex of these (flipped) adhered cells in the same focal plane. This set-up was thus used to image actin and NMII in the nuclear and perinuclear regions of adhered cells. We stained NMII monomers, however, single monomers might not have been detected due to their low fluorescence intensity. For suspended cells, no flipping of the samples was required, as the cortex images were taken directly.

**Figure 4:**
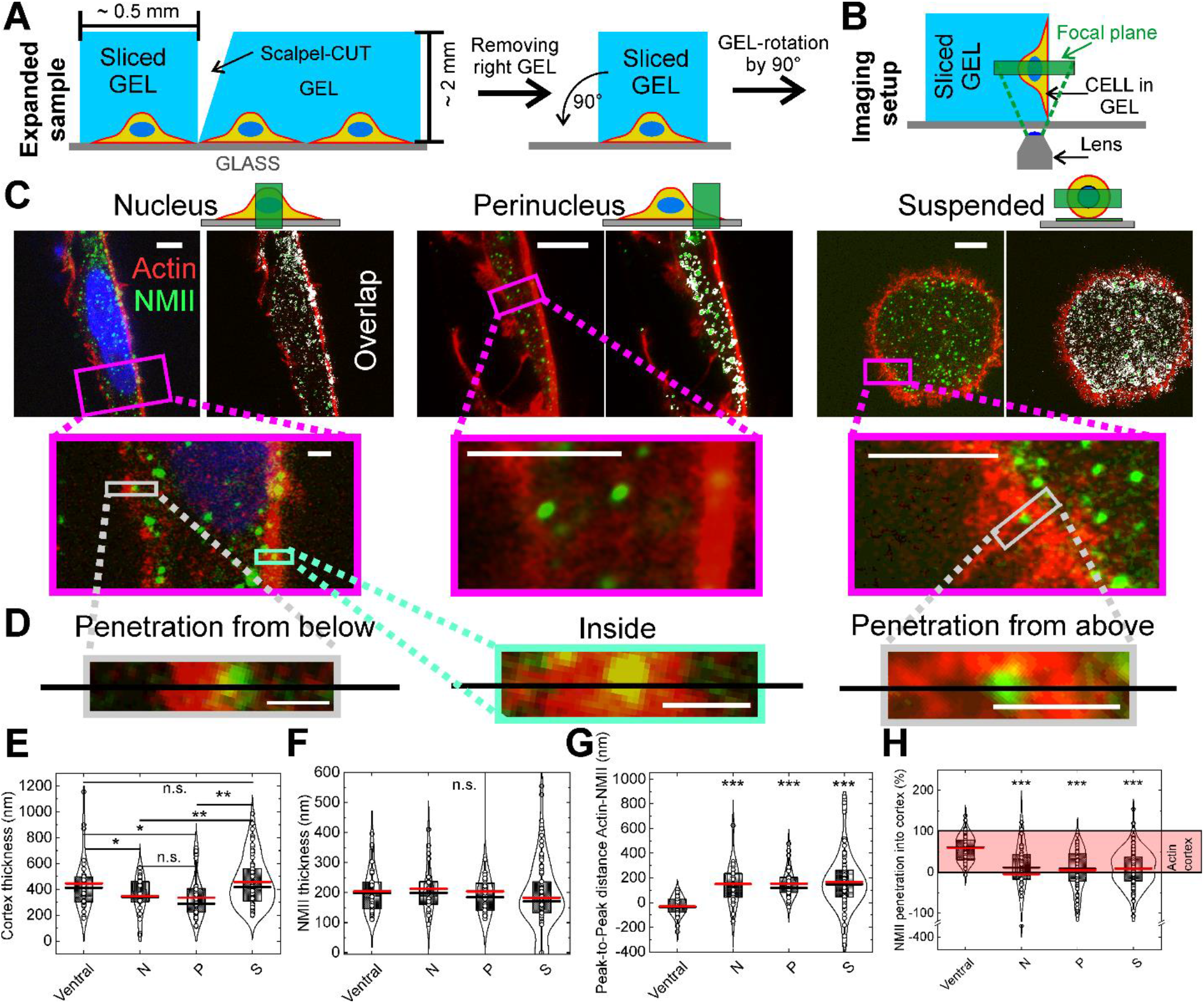
NMII minifilaments are not located within actin cortex of hTERT-RPE1 cells. Scheme of the preparation procedure for side-view imaging of adhered cells using expanded samples (**A**). Final imaging setup after gel (and hence cells) were rotated by 90° to enable side-view imaging (**B**). Side-view of expanded nuclear (scale bar: 2 μm, magnification scale bar: 500 nm) and perinuclear (scale bar: 1 μm, magnification scale bar: 500 nm) regions as well as suspended (scale bar: 1 μm, magnification scale bar: 250 nm) RPE1-cells imaged with Expansion microscopy in combination with Airy-Scan (**C**). Along the black lines the intensity profiles were measured (scale bars: identical to magnifications of (C)) (**D**). Analysis of intensity profiles leads to actin cortex thickness (**E**), NMII thickness/diameter (**F**), Peak-to-Peak distance between actin and NMII signals (**G**) and Penetration of NMII into the actin cortex (illustrated in light red) (**H**). Median values are marked in red, mean values in black. Star method is representing statistical Welch-corrected t-tests: n.s.: not significant, *: p<0.05, **: p<0.01, ***: p<0.001. Cell counts: ventral: n= 10, dorsal (nucleus, N): n= 6, dorsal (perinucleus, P): n= 10, suspended,S: n= 13. Numbers of total measurements: ventral: n_V_= 32, nucleus: n_N_= 21, perinucleus: n_P_= 32, suspended cells: n_S_= 78 (numbers are doubled for NMII penetration (G) due to upper and lower NMII penetration distance).

On the dorsal cell side, the NMII signals were only seen close to the actin cortex, and rarely within the actin cortex, independent of the NPS state (Figure 4C). In contrast, the NMII signal co-localized with the actin signal on the ventral cell side. Quantitative analysis showed that the peak signal of actin was not in the same position as the peak signal of NMII, except for the ventral side. In particular, this showed that the NMII peak signal was mainly on the inner side of the dorsal actin cortex (Figure S4A, B). However, there might have been nonbundled NMII in the actin cortex, but at levels below the resolution limit here. We further analyzed the intensity profiles to quantitatively extract the thicknesses of the actin and NMII layers, as well as their overlap (Figure S4B). For the ruffle-free areas, the actin cortex thickness was 300 nm in the adhered cells, and 400 nm in suspended cells (Figure 4E), with a thickness/ diameter of the NMII signals of ~200 ±50 nm, independent of the NPS state (Figure 4F). As the signal of single myosin protein is not expected to be detected in our experimental conditions, we identified these as NMII minifilaments [27–33]. The distance from the actin peak signals to these NMII minifilament peak signals was ~200 nm for all of the NPS states, although not for the ventral side of the cells, due the presence of NMII in stress fibers located at the ventral side of the cells (Figure 4G) which have been shown to penetrate the ventral actin cortex [34]. As both the actin cortex and NMII thicknesses can be >200 nm, we quantified the penetration of the NMII signals into the actin signal. This revealed that the NMII minifilaments penetrated the actin cortex by a mean depth of ~10% into the actin cortex thickness (Figure 4H).

The cortical mesh hole diameter ranged from 55 nm to 75 nm on average (Figure 2), and there was no indication of significantly larger MHA on the inner side of the actin cortex. Consequently, it is reasonable that NMII filaments with a length of 250 nm and a diameter of 100 nm [31, 33] would not enter the dorsal actin cortex. This would explain why no co-localization of actin and NMII signals was observed in the expansion microscopy images. Moreover, this supported our hypothesis that the absence of NMII minifilaments from the actin cortex leads to only minor influences on both actin cortex structure and cellular mechanics (for small indentations on short time scales). The cellular mechanics investigated with AFM and the lower actin t_1/2_ upon blebbistatin treatment are discussed further in the Supplementary Information.

To summarize here, we observed that NMII (minifilaments) were not located in the actin cortex, but were just beneath the actin layer on the dorsal side of the cells. This position allows only little interactions of the NMII minifilaments directly with the actin cortex, which explains the minor effect in MHA and E we observed after treatment with Blebbistatin and Calyculin. This separation of the actin layer from the NMII minifilament layer is in strong contrast to the ventral side of the cells, where NMII (minifilaments) are recruited during the formation of stress fibers and thereby fully embedded in the cortex [35, 36].

As the structure of actin is not only dependent on the actin itself and the NMII activity, but also on actin nucleation and elongation factors, we also investigated the effects of Arp2/3 and formins.

### Arp2/3 and formin influence the different cell adhesion states

The aspects of Arp2/3 and formin inhibition and localization are extensively discussed in the Supplementary Information. In brief, inhibition of Arp2/3 (using CK-666) and formin (using Smifh2) appeared to affect the DSM measures similarly for the actin cortex over the nucleus, while the effects of formin inhibition were stronger than Arp2/3 inhibition for that of the perinucleus (Figure 2). As Arp2/3 and formin show similar distributions in adhered cells, we concluded that both are highly relevant for the actin cortex over the nucleus, while formin is more relevant for the actin cortex of the perinucleus (Figure S5C-G). For suspended cells, again both Arp2/3 and formin were similarly distributed, with slightly increased levels for the actin cortex compared to the cytoplasm (Figure S5A, B). As there were not any large effects on the DSM measures for the CK-666 and Smifh2 treatments (Figure 2), we concluded that both Arp2/3 and formin have minor roles within the actin cortex of suspended cells, as long as there are no direct mechanical interactions, as further explained in the Supplementary Information. In summary, whilst Arp2/3 and formin are similarly distributed, both impact the actin cortex above the nucleus but formin is more relevant than Arp2/3 in the perinucleus and neither play a large role in suspended cells.

## Conclusions

The actin cortex is involved in a number of physiological phenomena, including processes that involve shape changes, such as cell polarization and migration. Therefore, it is important to characterize the actin cortex properties and their correlations upon cortical alterations. We found clear predictable correlations between actin dynamics, cortical structure and cellular mechanics (DSM) measures upon extrinsic alterations of the actin cortex of RPE1 cells, which show mesenchymal characteristics even though they are of epithelium origin. However, effects of these external perturbations are only partially correlated between different adhesion states. Interestingly, actin cortex alterations induced by the adhesion state transition per se (intrinsic alterations) lead to fundamentally different actin cortex behaviors, which emphasizes how cells – here with the example of the actin cortex – change during biological processes. In particular, we find that correlations between DSM measures of the actin cortex are different upon intrinsic alterations compared to extrinsically induced alterations. The most striking difference is that higher E is correlated with smaller Mesh Hole Area (MHA) for extrinsic alterations but with larger MHA for intrinsic adhesion state changes. In conclusion, extrinsic alterations provoke the mechanical homeostasis of cells, while intrinsic alterations accompany necessary requirements to fulfil function and enable further development.

We observed surprisingly minor effects on the structure and mechanics of the actin cortex upon inhibition or activation of key actin-regulating proteins including Arp2/3, formin and myosin. From our data, we predict that the main differences in the actin cortex between suspended cells and adherent cells is due to the down-regulation of the ROCK kinase in suspended cells. As well as the correlations of the adhesive states of the cortex over the nucleus, at the perinucleus and of suspended cells (NPS states) under intrinsic and extrinsic alterations, we were surprised to see minor effects on the structure and cellular mechanics upon myosin inhibition and activation. Our high-resolution imaging of myosin reveals that the NMII minifilaments are not – as believed to date – localized within the actin cortex, but underneath it. This structural separation of the actin and NMII layers can indeed explain a number of opposing results in the literature, such as the question how cellular mechanics alters under myosin inhibition. With our model, the answer to this question depends among other parameters on the amount of indentation of the cortex. If a particular method globally deforms whole cells or indent locally enough so that both the actin and the myosin layer are deformed, effects of myosin inhibition will be noticed. If the local deformation is little and will only deform the actin layer itself, no effects of myosin inhibition will be observed.

Taken together, we present here a comprehensive overview of actin cortex parameters and their correlations, which are in line with the vast quantity of data collected over decades by ourselves and others for particular actin cortex parameters. The discrepancies in the literature at present can be unified by our new proposition of the structure of the actin cortex. This defines the NMII minifilaments as forming their own layer beneath the actual actin cortex layer. We believe that this provides the basis for a new understanding and better control of the actin cortex and its behavior in living cells.

## Abbreviations

DSM: actin dynamics, cortical structure and cellular mechanics
NPS: The dorsal actin cortex over the nucleus, at the perinucleus, and of suspended cells.
Adhesion states: Cells which are adhered (N and P) or suspended (S). Although N and P are different regions of adhered cells, their fundamental differences (e.g., stress fibers) might reflect a different adhesion state.
Extrinsic (cortical alterations): Cortical alterations induced by cytoskeletal perturbants
Intrinsic (cortical alterations): Cortical alterations induced by adhesion state transitions (adhered versus suspended cells) or regional differences within the adhered cell state (N versus P)
MHA: <u>M</u>esh <u>H</u>ole <u>A</u>rea as “focus parameter” for the cortical structure.
t_1/2_: Actin half-time recovery of cortical actin as “focus parameter” for the cortical dynamics.
E: Stiffness of cells as “focus parameter” of the cortical mechanics.
NMII: Nonmuscle myosin II

## Acknowledgements

The authors thank **Johannes Rheinländer** (Department of Nanobiophysics and Medical Physics, Eberhard-Karls University Tübingen, Tübingen, Germany) for the Creep compliance analysis implementation and outstanding support in terms of discussions. Moreover, the authors thank **Ewa K. Paluch** (Department of Physiology, Development and Neuroscience, University of Cambridge, United Kingdom) for fruitful discussions and especially the discussions on whether nonmuscle myosin II is located within the actin cortex. The authors would also like to thank **Matthieu Piel** (Paris, France) for the hTERT-RPE1 cells stably expressing mCherry LifeAct. For financial support, the authors would like to thank the Leibniz Institute for New Materials (INM), Saarland University, the DFG (CRC 1027 (A10)) and the Max Planck School Matter to Life.

## Author contributions

DF, ET and FL designed and supervised the study

DF, KK, DV, CA, MP, LK and BG performed experiments and analyzed data (DF: SEM, AFM, Expansion Microscopy; KK: FRAP; DV: Immunofluorescence of Arp2/3 and Formin; CA: AFM; MP: AFM; LK: Immunofluorescence cell size; BMG: almar blue assays)

MS implemented FiNTA (Filament Network Tracing Algorithm)

AJ provided essential ideas and contributions within several discussions, and specifically for AFM RH provided essential ideas and contributions within several discussions, as well as outstanding theoretical expertise in the understanding of the actin cortex.

DF, KK, DV, ET and FL wrote and revised the manuscript

KK, AJ and RH revised the manuscript

## Materials and Methods

For general details of the sources of the materials used in this study, see also Table 1.

**Table 1.**
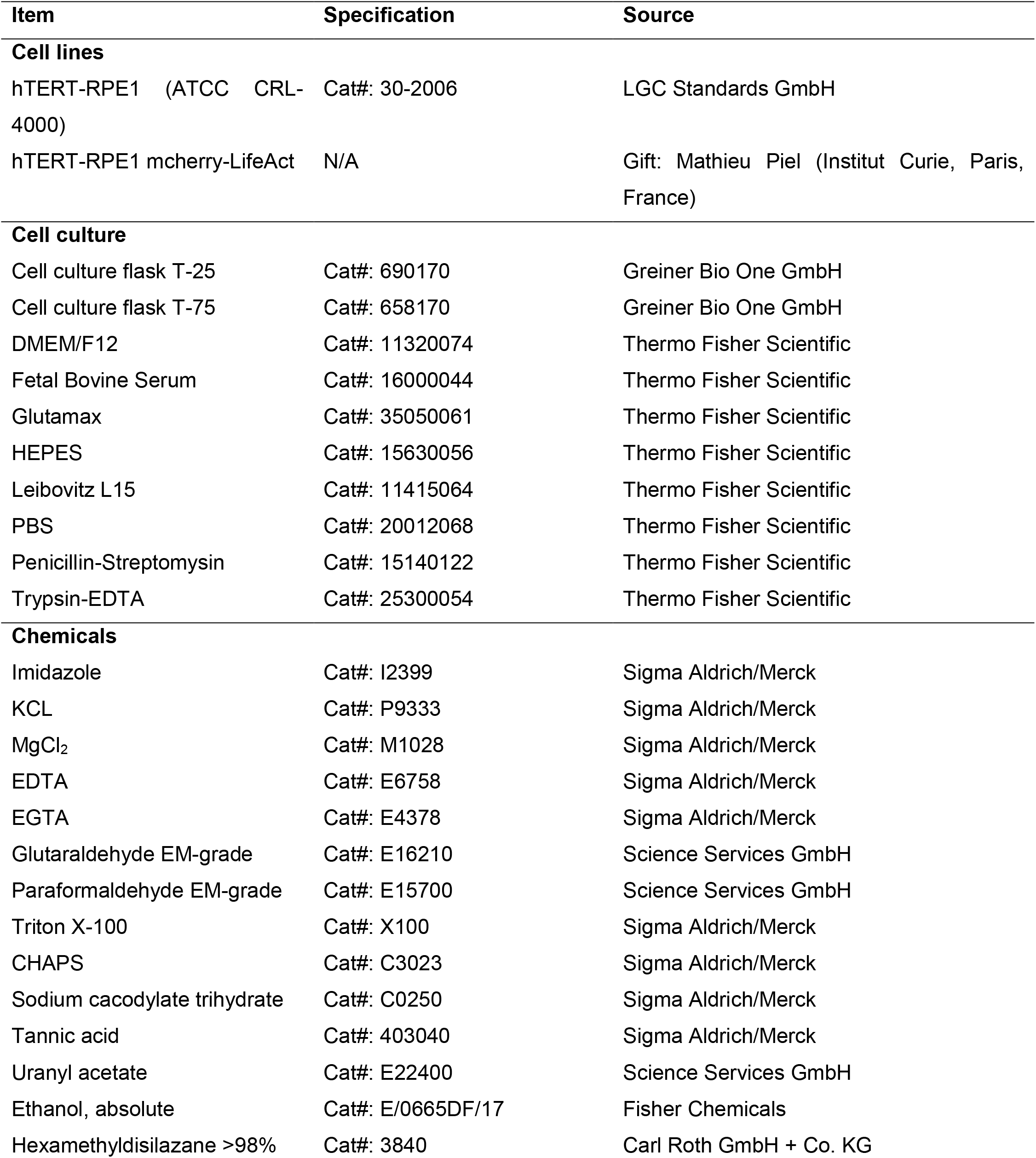
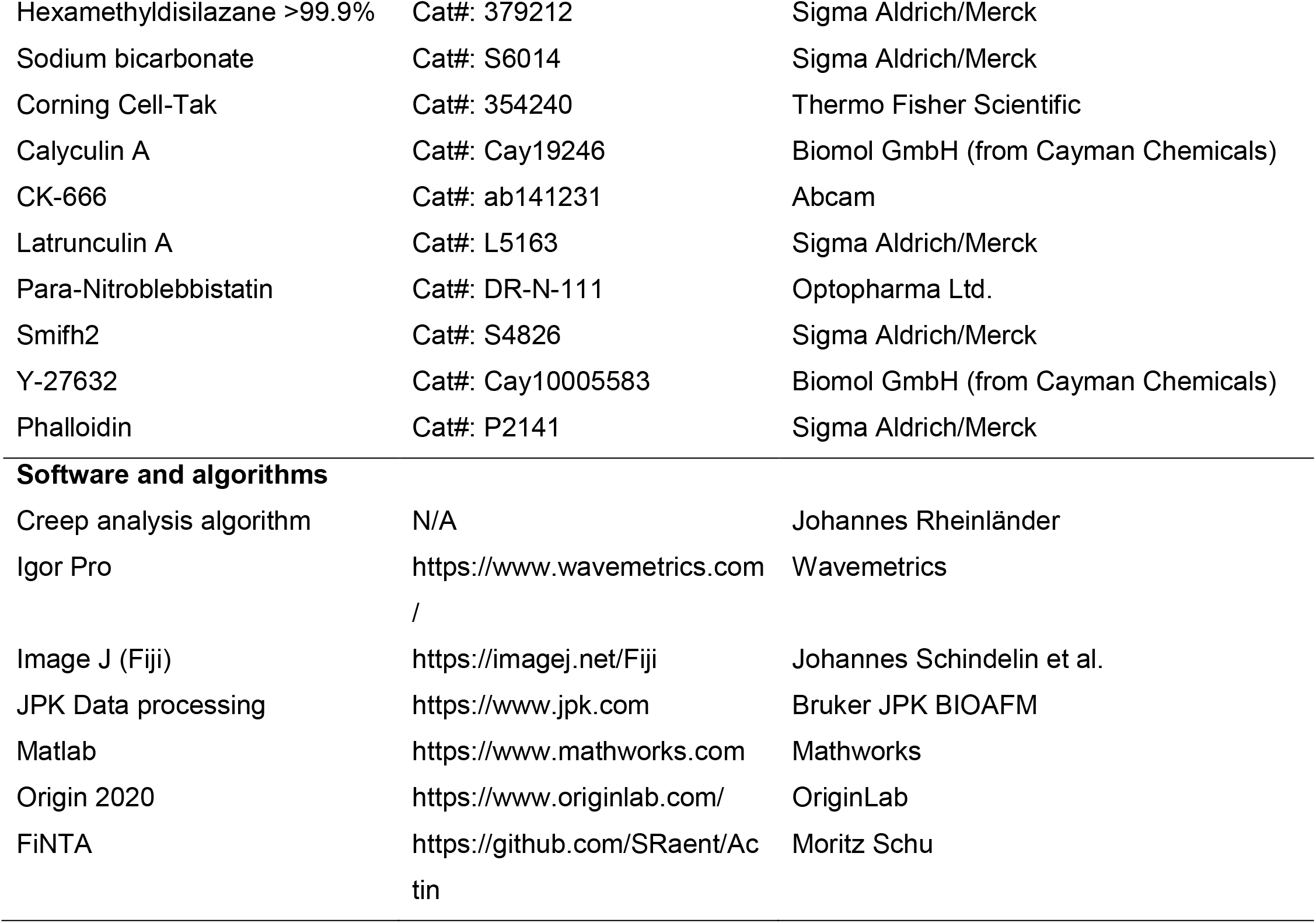

### hTERT RPE1 cell culture

Telomerase-immortalized (hTERT-) retina pigmented epithelium (RPE1) cells (LGC Standards GmbH, Germany) were cultured in DMEM/F12 medium (Thermo Fisher Scientific, Germany) supplemented with 10% fetal bovine serum (Thermo Fisher Scientific, Germany), 1% penicillin-streptomysin (Thermo Fisher Scientific, Germany), and 1% Glutamax (Thermo Fisher Scientific, Germany) in cell culture flasks (Greiner Bio-one GmbH, Austria). The culture conditions were 95% air and 5% CO_2_ at 37 °C. Cells were passaged every 3 days, and preserved for a maximum of 15 passages.

For investigation of cells in suspension (suspended cells) and for passaging, the cells were detached using trypsin (Thermo Fisher Scientific, Germany) at 37 °C under 5% CO_2_ for 3-5 min. The cell suspensions were centrifuged at 300× g, and the supernatants were exchanged for culture medium (with cytoskeletal agents later included as necessary). During the incubations of cell suspensions, the reaction tubes were constantly rotated at 360°/min, to avoid cell adhesion to the tubes. For further investigations of the adhered and suspended cells with AFM and FRAP, the culture medium included 25 mM HEPES (Thermo Fisher Scientific, Germany).

### Genetic modifications

To conduct FRAP measurements, the cells were transfected using X-tremeGENE 9 DNA Transfection Reagent (Roche, Germany) with a plasmid encoding γ-actin N-terminally tagged with green fluorescent protein (GFP). The plasmid was a kind present from A. Simiczyjew (University Wroclawski, Poland). The transfected cells were selected using 400 μg/mL G418 for 5 days. Subsequently, the transfected cells were cultured with 200 μg/mL G418.

### Cytoskeletal perturbations

Unless indicated otherwise, the cells were treated with the following cytoskeletal agents at 37 °C: 20 μM para-nitroblebbistatin for 30 min (NMII ATPase inhibitor; Optopharma Ltd., Hungary); 10 μM Y-27632 for 30 min (ROCK-inhibitor; Biomol GmbH, Germany); 1 nM calyculin A for 30 min (protein phosphatase PP1 and PP2A inhibitor; Cayman Chemicals, MI, USA); 100 μM CK-666 for 30 min (Arp2/3 inhibitor; Abcam, UK); 10 μM Smifh2 for 30 min (formin FH2 domain inhibitor; Sigma-Aldrich, Germany); and 0.1 μM latrunculin A for 10 min (actin polymerization inhibitor; Sigma-Aldrich, Germany). No cytoskeletal perturbant treatment of hTERT-RPE1 cells led to significant cell death (Figure S9C, D).

### Scanning electron microscopy

The preparation of the hTERT-RPE1-cells for SEM was as described by [37–39], with minor modifications. SEM micrographs were obtained (Quanta 400; Thermo Fisher Scientific, Germany) at 5 kV, spot size 2 in high vacuum mode. The resulting images were analyzed using the filament network tracing algorithm FiNTA [40]. For quantification of the filament length and network connectivity, the unification distance was set to 2. Furthermore, the break-off angle to determine the filament length was set to 0.3 rad (~17°). Implemented intensity thresholding was used to exclude very bright actin cortex regions (e.g., most frequently ruffles in suspended cells) and relatively dark actin cortex regions (e.g., most frequently background filaments that did not belong to the same focal plane as the cortical region of interest). Therefore, the grayscale intensity of interest for most images was between 30 and 230.

### Preparation of adhered cells for scanning electron microscopy

For SEM, the cells were plated overnight onto glass coverslips (diameter, 25 mm; Thermo Fisher Scientific, Germany) at 37 °C for at least 24 h. The following procedure was performed at room temperature, unless indicated otherwise. After optional treatments with the cytoskeletal perturbants as described above (see **Cytoskeletal perturbations**), the cells were rinsed three times with serum-free Leibovitz L15 medium (Thermo Fisher Scientific, Germany). Without washing, the L15 was removed and extraction solution #1 was added for 5 min, which contained cytoskeleton buffer (50 mM imidazole, pH 6.8, 50 mM KCl, 0.5 mM MgCl_2_, 0.1 mM EDTA, 1 mM EGTA) with 10 μM phalloidin (Amanita phalloides; Sigma Aldrich, Germany), 0.25% glutaraldehyde (EM grade; Science Service GmbH, Germany) and 0.5% Triton X-100 (Sigma Aldrich, Germany). Without washing the cells, extraction solution #1 was replaced by extraction solution #2 (2% Triton X-100, 1% CHAPS hydrate; Sigma Aldrich, Germany) for 5 min. Both procedures are intended to remove the cellular membranes and pre-fix the actin network. After rinsing three times with cytoskeleton buffer, the solutions were replaced by the fixing solution overnight (2% glutaraldehyde, 2% paraformaldehyde [EM grade; Science Service GmbH, Germany] in 100 mM aqueous sodium cacodylate buffer [Sigma Aldrich, Germany]). Without rinsing the cells, the fixing solution was replaced by 0.1% aqueous tannic acid (Sigma Aldrich, Germany) for 20 min. After rinsing three times with MilliQ water, a final concentration of 0.2% aqueous uranyl acetate (Science Service GmbH, Germany) was added for 20 min. After rinsing three times with MilliQ water, ethanol (Fisher Chemicals, Germany) was added gradually using a 60-mL syringe pump (Harvard Apparatus, United states) as two cycles: first, the water was exchanged for 50% aqueous ethanol over 45 min, and then the 50:50 ethanol:water solution was exchanged with 100% ethanol over 45 min. After rinsing three times with 100% ethanol, where the samples were dried over a molecular sieve for 15 min after every rinse, 1,1,1,3,3,3-hexamethyldisilazane (HMDS; Carl Roth GmbH and Co. KG, Germany) was added successively at 5%, 10%, 20%, 30%, 40%, 50%, 60%, 70%, 80%, 90%, 95%, >98%, for 5 min at each concentration. After three incubations for 10 min each with 99.9% HMDS reagent plus (Sigma Aldrich, Germany), the HMDS was evaporated off under a chemical hood. Immediately after the final drying, the samples were sputtered with 6-7 nm platinum (Coater: Model 681; Gatan, USA).

### Preparation of suspended cells for scanning electron microscopy

Coverslips were coated Cell-Tak (Corning Cell-Tak; Fisher Scientific, Germany) in 0.1 M sodium bicarbonate (Sigma Aldrich, Germany) for 20 min, as described in the Basic Adsorption Coating Protocol (Fisher Scientific, Germany), followed by three washes with phosphate-buffered saline (PBS; Thermo Fisher Scientific, Germany) for 10 min each.

After cell detachment and optional incubation with the cytoskeletal perturbants (see above), the cells were rinsed three times with L15, using centrifugation (see above). Without washing the cells, the L15 was removed and replaced by extraction solution #2 supplemented with 10 μM phalloidin, for 9 min. After removal of the PBS from the coverslips, the cell extraction suspension was added to Cell-Tak–coated coverslips for 1 min. The further steps were identical to the SEM preparation for adhered cells (see above).

### Preparation for atomic force microscopy

The local mechanical properties of adhered and suspended cells were determined using an AFM system (Nanowizard 3; JPK Instruments/Bruker, Germany) mounted on an optical microscope (Eclipse Ti-U, Nikon, Minato, prefecture Tokyo, Japan), equipped with PlanFluor 40x, NA 0.6, Ph2 objective (Nikon). Cantilevers (MLCT-C, Bruker, France) with a pyramidal indenter were used (nominal tip radius, 20 nm; nominal spring constant, 0.01 N/m; nominal resonant frequency, 7 kHz). The setpoint force was set to 400 pN to maintain indentations of the order of the cortical thickness (see Figure S11). The approach velocity was set to 5 μm/s.

### Creep investigations

Creep investigations are based on the studies of Hecht et al. (2015) [41], who described the theoretical background in more detail. In brief, the cells were defined according to a power-law material. The investigation was based on the force-clamp method. Here, the cantilever was approached at 5 μm/s until a clamp force of 400 pN was reached. This force was kept constant for 1 s, while a feedback loop controlled the z-position of the cantilever. During this force-clamp period, the indentation of the cantilever into the cell was increasing to keep the force constant.

The creep compliance equation shown by Equation (1) was used for power-law materials.

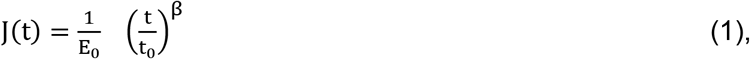

where E_0_ is the scaling parameter, or stiffness, and β is the fluidity (0 < β < 1). When β = 0, the material behaves as an elastic solid, when β = 1, the material behaves as a viscous Newtonian fluid.

For comparisons, the approach segment was used to extract the Young’s modulus, by the classic Hertz-Sneddon approximation [42, 43]. As this model treats a cell as a purely elastic material, the viscoelastic behavior of the cell was not taken into account. Nonetheless, using a high approach velocity (e.g., 5 μm/s), the Hertz-Sneddon approximation can be used by focusing on the purely elastic characteristics of a cell.

### Fluorescence recovery after photobleaching

For all of the FRAP measurements a confocal laser-scanning microscope was used (LSM 880) equipped with an Airyscan detector (Zeiss, Germany). The cells were visualized under an oil immersion objective (63×, 1,4 NA), and images were acquired with a 488-nm laser diode and a dwell time of 3 μs per pixel, at 10% laser power. For bleaching, a circular region of interest with a diameter of 2 μm was illuminated using a 405-nm laser diode at 20% laser power with a dwell time of 177 μs per pixel. The total acquisition length was 150 s, with a frame rate of 4 fps. Image analysis, imaging-induced photobleaching, and data extraction was performed using a custom written MATLAB script. Equation (2) was used to fit the data obtained.

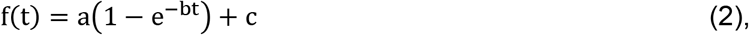

where a, b, and c are the free parameters that refer to the plateau value, turnover rate, and intensity offset, respectively. For the presented discussion, only the plateau a and the derived half-time recovery, 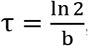, were considered.

### Sample preparation for FRAP measurements

For FRAP measurements with adhered cells, the cells were first detached using trypsin, and counted. Subsequently, the cells were plated on circular coverslips (diameter, 25 mm; thickness, 0.17 mm) inside each well of sterile six-well plates (40,000 cells/well), overnight with DMEM/F12. An hour prior to the experiments, metallic cell culture chambers (SKE Research Equipment) were cleaned with ethanol and submerged in DMEM/F12 with 25 mM HEPES for at least 30 min. For the perturbant treatments, this medium was additionally supplemented with the perturbant. The glass coverslips with cells were then rinsed with Dulbecco’s PBS (DPBS) and positioned in the tight fitting of the metallic cell culture chamber. Then, 2 mL DMEM/F12 with HEPES was added to each chamber, and a clean glass coverslip (diameter, 25 mm; thickness, 0.17 mm) was put on top of the liquid and gently pressed down to close the chamber. The residual liquid was then aspirated off.

For the FRAP measurements on suspended cells, custom build chambers were used. A piece of gel film (PF Gel-Film X4; Gel-Pak, USA) with a thickness of 6.5 mil (165 μm) was put on a folded piece of Parafilm, and multiple holes were made in the PF Gel-Film using a Harris unicore punch (diameter, 2.5 mm; Merck, Darmstadt). Then the piece of PF Gel-Film was put into the center of a sterile cell culture dish (FluoroDish FD35-100; Ibidi, Germany). The FluoroDish with the PF Gel-Film together with a clean glass coverslip (diameter, 22 mm; thickness, 0.17 mm) was then put in a plasma cleaner (Plasma Lab System Atto; Diener Electronic GmbH and Co., KG, Germany) and activated for 3 min. After this activation, 20 μL poly-L-lysine grafted polyethyleneglycol copolymer (PLL-g-PEG) solution was added to each punched hole of the PF Gel-Film, and a piece of Parafilm was put on top and gently pressed down. If any air pockets were formed, the piece of Parafilm was lifted, and the process was repeated. Then, 40 μL PLL-g-PEG solution was pipetted onto a piece of Parafilm, and the glass coverslip was put on the droplet and gently pressed down. The PLL-g-PEG was allowed to bind to the glass surface for at least 1 h.

Then, 1 h prior to the experiments, the cells were detached using trypsin and counted. A total of 1,000,000 cells was put into a 1.5-mL microcentrifuge tube (Eppendorf), and centrifuged at 150× g for 3 min in a tabletop centrifuge. The supernatant was aspirated off, and replaced with 1 mL fresh DMEM/F12 with 25 mM HEPES, and put on a shaker pre-installed in an incubator, and shaken for 60 min, to keep the cells in suspension. Then, 30 min prior to the experiment, the parafilm inside the FluoroDish was removed, the dish was rinsed three times with sterile DPBS, and 2 mL DMEM/F12 with 25 mM HEPES was added to let the PF Gel-Film absorb the liquid and the supplements it contained. Immediately prior to the experiment, the liquid in the FluoroDish was aspirated, and the FluoroDish was rinsed once with DPBS. The microcentrifuge tube containing the cells was then taken out of the incubator, and 10 μL of the cell suspension was pipetted directly into each of the punched holes. The glass coverslip was then lifted from the Parafilm and rinsed once with DPBS. Any residual DPBS droplets were aspirated off, and the glass coverslip was put on top of the PF Gel-Film with the PLL-g-PEG coated side facing downwards.

For the perturbant treatments, the microcentrifuge tube containing the cells was taken out of the incubator after 30 min, centrifuged once at 150× g for 2 min, and the supernatant was aspirated and replaced with fresh DMEM/F12 containing 25 mM HEPES and the relevant perturbant at the concentration required (see above). The DMEM/F12 with 25 mM HEPES added to the FluoroDish was also supplemented with the relevant perturbant at the concentration required.

### Immunostaining and fluorescence microscopy

#### Immunostaining

Arp2/3 and formin: The day before the immunostaining, cells were plated on top of the coverslip and incubated overnight at 37 °C and under 5% CO_2_, to allow them to adhere. CellTak (Corning Cell-Tak; Fisher Scientific, Germany) was diluted in 0.1 M NaCO_3_ and 1 M NaOH, in the proportion given in manufacturer instructions, and put on top of the coverslips for 20 min. The coverslips were then rinsed three times with phosphate-buffered saline (PBS; Thermo Fisher Scientific, Germany) and stored at 4 °C overnight.

The suspended cells were trypsinized and agitated for 30 min in HEPES-supplemented DMEM/F12 medium at 37 °C and under 5% CO_2_. The cells were then incubated with 3% bovine serum albumin (BSA) for 1 h, to minimize nonspecific binding of the primary antibodies. The primary antibodies used for the immunostaining were mouse monoclonal antibodies (Arp3 [A-1], sc-48344; formin 2 [C-3], sc-376787; Santa Cruz Biotechnology, Dallas, Texas, USA). The primary antibodies were diluted 1:200 in 3% BSA and left in contact with the cells for 1 h at room temperature, with the cells then rinsed three times with BSA. The Alexa Fluor 488 donkey anti-mouse secondary antibody (Abcam, Cambridge, UK) and phalloidin conjugated with tetramethyl rhodamine B isothiocyanate (Sigma Aldrich, St Louis, MO, USA), were diluted 1:1000 in 3% BSA, and left in contact with the cells for 1 h at room temperature. After this staining, the cells were rinsed three times in MilliQ water. Then the glass coverslips were mounted on glass slides using mounting medium containing DAPI to stain the nuclei (Fluoromount-G, with DAPI; Thermo Fisher Scientific, Waltham, MA, USA). The mounted glass slides were kept overnight at room temperature in the darkness.

#### Fluorescence microscopy

Arp2/3 and formin: Epi-fluorescent images were acquired using an inverted microscope (Ti-Eclipse; Nikon, Melville, NY, USA) equipped with a monochrome CCD camera of 6.45 μm pixel size (DR328; Andor Technology, Belfast, UK) at a magnification of 60× and a binning of 2×2. The image analysis was carried out using the plugin Radial Profile Angle of the Image J software (Fiji), with a circle of 43 μm centered on the nucleus.

#### Expansion microscopy

Expansion microscopy is a technique that is used to increase the spatial resolution of fluorescence microscopy images by a factor of 4.5. In expansion microscopy, the cells are essentially permeated by a hydrogel, which is then swollen by the addition of water. The preparation followed the protocol for cultured cells of Chen et al. (2015) [26]. In brief, after fixing of the hTERT-RPE1 cells stably expressing mCherry-lifeact with 0.5% paraformaldehyde, and immunostaining for NMII-MLC (see above), the samples were anchored with Acyloyl-X for 3 h. Samples were polymerized with a hydrogel for 1 h at 37 °C. After digestion (including ProteinaseK), water was added to the samples. In this way, the sample size was physically increased by a factor of 4.5 in all spatial directions.

To acquire high resolution images of the ventral and dorsal sides of the actin cortex of the cells, the slices of cells embedded in gel were flipped through 90°, directly onto a glass dish (Figure 4A, B).

### Statistics

#### Minimum-maximum normalization

All of the DSM measures for the focus parameters and the NPS states were assigned to different scales with minor overlap (Figure S1A-C). For direct comparisons (correlations) the means of the data were normalized using the proportion of maximum scaling (POMS; so-called minimum/maximum) method. Here, the maximum value of each focus parameter per NPS state was set to 1 and the minimum to 0, and the remaining values were re-calculated from these two extreme values. This method allowed direct comparisons between the DSM measures without changing the Pearson R correlation coefficient between them, compared to un-normalized data. Since we investigated correlations with and without Y-27632, data were re-normalized, if any Y-27632 measure was equal to 1 (see Figure S1D-I).

#### Data presentation and statistical tests

All of the data are presented in violin plots for direct visualization of the data distributions. In all Figures, the red horizontal lines represent the means of the distributions, and the black horizontal lines represent the medians of the distributions. Mann–Whitney U-tests were performed for statistical analysis between the investigated conditions. Here, p <0.05 defined statistical significance. Unless specified otherwise, statistical significance refers always to the control condition (untreated hTERT-RPE1 cells). Color-coded significance indicators are exclusively used to compare control conditions for the NPS states (green), as well as just for the perinucleus region and suspended cells (violet). The use of n.s. or the absence of any statistical indication defines conditions that were not significantly different from each other.

#### Cell viability with the Alamar Blue assay

The hTERT-RPE-1 cells were seeded into 96-well plates (2,000 cells/well). After 24 h, the different perturbants where added to the following final concentrations: 20 μM blebbistatin; 100 nM latrunculin A; 10 μM Smfih2; 1 nM calyculin; 100 μM CK666; 10 μM Y-27632; or CK666+Smifh2. All of the treatments were carried out for 30 min, except from the latruncuin A, which was incubated for 10 min. After the treatments, the cells were rinsed. Then the cells were incubated in fresh medium for 1 h, and 20 μL of Alamar Blue (Invitrogen) reagent was added to each well in 180 μL medium. The cells were incubated for 16 h at 37 °C in a CO_2_ incubator, to allow the Alamar Blue to be metabolized. The absorbance of the samples was measured at 540 nm (using 620 nm as a reference wavelength) using a microplate reader (Infinite200, TECAN branch, Switzerland). The assay was performed using four replicate wells for each perturbant tested.

