## Supplementary information for "The role of actin and myosin II in the cell cortex of adhered and suspended cells"

**Short title:** Cell cortex of adhered and suspended cells

D.A.D. Flormann[1,2]*, K.H. Kaub[1,2,3], D.Vesperini[1], M. Schu[1], C. Anton[1], M.O. Pohland[1], L. Kainka[1], G. Montalvo Bereau[1], A. Janshoff[3] , R.J. Hawkins[4], E. Terriac[1,2], F. Lautenschläger[1,5]*

[1] Department of Physics, Saarland University, Saarbrücken, Germany

[2] Leibniz Institute for New Materials, Saarbrücken, Germany

[3] Department of Biophysical Chemistry, Georg-August-University, Göttingen, Germany

[4] Department of Physics and Astronomy, University of Sheffield, Sheffield, United Kingdom

[5] Centre for Biophysics, Saarland University, Saarbrücken, Germany

In the present study, the activities were altered for the three main components considered here: nonmuscle myosin II (NMII), Arp2/3, and formins. This was achieved using small inhibitory molecules with adhered and suspended hTERT-immortalized (hTERT)-RPE1 cells. NMII was directly interfered with using blebbistatin and calyculin A (a phosphatase inhibitor that actives NMII via its actions on PP1 and PP2A phosphatases). Furthermore, the NMII activator Rho-associated protein kinase (ROCK) was inhibited by treating the cells with the ROCK inhibitor Y27632. Also, CK-666 was used to inhibit Arp2/3, and finally, formins activity was inhibited with the inhibitor of formins-homology-2 (FH2) domain, Smifh2.

The resulting changes in the actin dynamics, the actin cortex structure, and the cellular mechanics (defined here as the dynamics, structure and mechanics [DSM] measures) under these various cell treatments were analyzed using fluorescence recovery after photobleaching (FRAP), scanning electron microscopy (SEM), and atomic force microscopy (AFM), respectively. These allowed quantification of the correlations between the separate components of the DSM measures. Secondly, the same dataset was used to investigate the mechanical and dynamic characteristics of adhered and suspended cells, in terms of changes to the structure of the actin cortex induced by interference with the cytoskeleton, and their correlations with cell adhesion state. Thirdly, we examined the effects on NMII of the loss of NMII minifilaments within the actin cortex. Finally, we assessed the regional and cell state differences upon Arp2/3 and formin inhibition, to better define their relative roles in adhered and suspended cells.

Here, we present additional data and discuss potential hypotheses that are not required to follow the main core of this study, but which we believe will be helpful for anybody working on the actin cortex.

**DSM measure correlations: Extrinsic alterations to the actin cortex unravel clear correlations between actin dynamics, structure, and mechanics.**

As shown in Figure S1D-F, there was separation between the NMII-targeted perturbants (including controls), as indicated by the blue background, and the actin-targeted perturbants, as indicated by the red background. Although calyculin A is known to target NMII, for suspended cells, the data here were in the region of the parameter space associated with actin modifications. Also of note, Y-27632 is a ROCK inhibitor that has severe downstream effects on both NMII and actin, and therefore the correlations here were examined with and without addition of Y-27632 (i.e., normalization with Y-27632, and also separate re-normalization without Y-27632). The separation between the NMII-targeted and actin-targeted perturbants were independent of ROCK inhibition (Figure S1G-I).

Figure 3D-E shows the parameter space diagrams for each of the DSM measures, to compare each of the NPS states with each other. The analogous POMS normalization and Pearson R correlation analysis were used, as seen for Figure 3A-C. Here, the actin half-time recovery ($t_{1/2}$) showed positive correlation between the NPS states (Figure 3D). *This implies that this focus parameter is independent of the cell adhesion state, which supports the idea that our chosen perturbations affect the actin dynamics in a similar manner in each of these states.*

Moreover, the actin cortex mechanics were positively correlated between the NPS states (Figure 3F), although the correlation between the perinucleus and suspended cells relied on ROCK inhibition. *This latter is mainly a result of the lack of correlation of cell stiffness, E, between the perinucleus of the adhered cells and suspended cells. This might originate from stress fibers or actin bundles in the perinucleus of adhered cells. Stress fibers and actin bundles are known to be under high tension compared to the actin cortex, which is localized over the nucleus of adhered cells, as well as over the nucleus of suspended cells, where stress fibers are absent [1]. Additionally, and from a purely geometric perspective, the cortex curvature increases from the perinucleus to over the nucleus, and further for suspended cells, which is indirectly associated with lower actin cortex tension [1-4]. When ROCK was extrinsically down-regulated, the mechanics of the adhered cells were strongly correlated to the mechanics of suspended cells (Figure 4 F). This supports ROCK kinase activity as crucially different for adhered cells compared to suspended cells, as has been suggested in the literature [5]. This strongly suggests that the underlying biological processes upon adhesion state transitions cannot be explained by the intuitive correlations presented in Figure 3A-C.*

Further, the actin cortex structure was positively correlated only within the adhered cells (Figure 4E). Neither the over the nucleas nor the perinuclear regions showed obvious correlations with the structures of suspended cells (Figure S3H, K). Consequently, the actin cortex structure of suspended cells reacts differently to that of adhered cells upon extrinsic actin cortex alterations.

As the nuclear and perinuclear regions of the adhered cells were positively correlated (see Figure 3D-F), the basic datasets for the two types of adhered cells were also combined and averaged for each condition (Figure S3M-O). Here, for the regional dependent correlations, the dynamics correlation for $t_{1/2}$ was positive between the adhered and suspended cells (Figure 4 M). For the structure, for mesh hole area (MHA), there was no correlation between adhered and suspended cells (Figure 4N). For the mechanics, the correlation for *E* was only seen when the cells were treated with Y-27632 (Figure 4O).

*Consequently, the actin dynamics correlations between the NPS states are intuitively expected to lead to correlations for the structure of the actin cortex between adhered and suspended cells. However, we observed no correlation between adhered and suspended cells for this property (Figure 3E). We hypothesize that this lack of correlation is induced by the underlying biological processes, as discussed further in the next section.*

**Intrinsic actin cortex alterations induced by cell adhesion state transitions reveal the underlying biological processes**

Counter-intuitively, mechanical correlations between adhered and suspended cells do not require structural correlations between the adhered and suspended cells (Figure 4G, J, M). Up to this point, we have seen that the DSM measures were correlated within the NPS states, and that specific DSM measures were correlated across the NPS states. To show this, the actin cortex was altered extrinsically by cytoskeleton perturbants when the cells were both adhered and suspended. In addition, the actin cortex is intrinsically different under each of the given NPS states. For simplicity, we initially focus in the following on the untreated cells only (Figure 2, controls): in contrast to the extrinsically induced correlations in Figure 3, the intrinsic alterations of the actin cortex primarily lead to the opposite correlations of the DSM measures compared to those within each of the NPS states (Figure S2).

This observation has the following implications for intrinsic alterations to the actin cortex in the untreated cells: first, and contrary to the extrinsic actin cortex alterations, MHA is negatively correlated with $E$. Secondly, $t_{1/2}$ is correlated with MHA, as expected, except for the actin cortex of suspended cells compared to that over the nucleus. Thirdly, $t_{1/2}$ is correlated with $E$ as expected for suspended cells compared to over the nucleus for adhered cells. We hypothesize here that the concept that increased actin half-time recovery leads to increased MHA and therefore to decreased cell stiffness is only correct for the extrinsic alterations of the actin cortex within each of the NPS states. In contrast, this is not correct if the actin cortex alterations are induced intrinsically (i.e., via adhesion state transition).

We believe that these data can be explained on the basis that the adhered cells are under tension and will therefore be stiffer, such that MHA is relatively ‘stretched’ compared to the more ‘relaxed’ cells when in suspension (Figure S9E, S10). Additionally, there are two particular actin cortex characteristics that are of importance here: first, compared to the actin cortex over the nucleus, the perinucleus is a thin cell region with denser fiber packing under higher tension, which can be expected to influence the actin cortex mechanics, as explained below. Secondly, a decrease in ROCK activity in suspended cells significantly changes the architecture of the actin cortex.

***The mechanics of the perinucleus***

The mechanics of the actin cortex in the perinucleus differs greatly from that over the nucleus for two main aspects. First, the cellular stiffness of the perinucleus is higher than that over the nucleus, as we and others have reported previously [1, 6]. Additionally, at 6 µm in height, the nuclear cell region is approximately 3.5-fold higher than the perinuclear region, at 1.7 µm [7]. Here, we measured the cortex thickness as ~300 nm at the dorsal side of the perinucleus, and 400 nm at the ventral side of the perinucleus (Figure 5). Consequently, a gap of only ~1 µm remains to contain all of the other cellular elements within the perinucleus, such as the microtubules and intermediate filaments. Therefore, we assume the need for dense packing of the filaments and cytoskeletal elements within the perinucleus. As the indentations of the cantilever used for the stiffness measures are in the range of 200 nm, substrate effects are assumed to be negligible. Altogether, therefore, we hypothesize that the increased stiffness is due to the dense packing of the actin and other cellular elements that are located in this cell region. This implies that the actin cortex has very limited space to deform upon cantilever indentation, and consequently, that the cell stiffness is increased.

Secondly, traction force microscopy investigations have shown maximal forces at cell edges and protrusions, which are transmitted along a decreasing force gradient via the stress fibers towards the nucleus [1]. High actin tension is known to lead to increased stiffness, as the deformability of the cells is reduced. Although there are stress fibers on the dorsal cell side over the nucleus, we have rarely observed dorsal stress fibers over the nucleus in SEM investigations.

In conclusion here, the dense fiber packing of the actin cortex and below it, combined with higher tension in the perinucleus compared to over the nucleus, would explain the increased cell stiffness of the perinucleus compared to over the nucleus. Interestingly, these high tensions appear to lead to increased MHA (Figure 2A). Intuitively, a stretched network (i.e., under high tension) will lead to both increased stiffness and increased MHA, which is consistent with our observations in the perinuclear region.

Consequently, and intuitively, the high actin tension in combination with the dense fiber packing appears to influence the cellular mechanics more than the actin cortex structure *per se*. This underlines a fundamentally different causal chain compared to the D (actin dynamics) to S (cortical structure) to M (cellular mechanics) progression that we suggested initially that was based on our observations of effects of extrinsic modifications.

***The structure of suspended cells***

For suspended cells, both MHA and *E* of the untreated cells (i.e., controls) were lower compared to those for the adhered cells (Figures 2, S2A). In contrast, with the extrinsic actin cortex alterations via the cytoskeleton perturbants, the decrease in MHA correlated with the increase in

*E* within the NPS states (Figure 3). We hypothesize that this counter-intuitive trend can be explained according to the particular characteristics of suspended cells *per se*: compared to adhered cells, suspended cells show natural down-regulation of ROCK and low actin cortex tension [2, 5].

ROCK is a multipotent kinase that affects NMII, the ezrin–radixin–moesin complex, and adducin, and activates phospho-cofilin [8, 9]. In further considerations here, we would exclude NMII, the ezrin–radixin–moesin complex, and adducin as potential candidates that are influenced downstream of ROCK. We have shown here that NMII alterations have little effect on both actin cortex structure and cellular mechanics, and it appears that inhibition of NMII via ROCK has little effect compared to the other proteins involved here. Moreover, the ezrin–radixin–moesin complex might have an important role in actin cortex structure via a postulated feedback loop with Rho [10, 11], although the level of any potential direct influence of the ezrin–radixin–moesin complex on the actin cortex structure appears to remain an open question. Therefore, we are not going to attribute the structural actin cortex changes to the ezrin–radixin–moesin complex directly. In contrast, adducin appears to have a strong influence on the actin cortex structure [12], although depletion of the actin-binding LIM protein 1 (abLIM1) leads to relatively long bundles of actin fibers [13]. As suspended cells showed short, mainly unbundled, actin filaments, adducin would appear to have a minor role here.

Consequently, we hypothesize that increased activity of phospho-cofilin has the major role in the actin cortex of suspended cells. Cofilin is known to sever actin filaments, and therefore the 'fluidize' the actin cortex [14, 15]. We indeed observed increasing fluidity by the Y-27632 treatment for the NPS states, as well as an intrinsic increased fluidity in the transition from the perinucleus to suspended cells (Figure 2). Moreover, the severing of actin filaments is shown by the significantly shorter actin filaments in suspended cells compared to the over the nucleus and the perinuclear regions (Figure 2B). Additionally, inhibition of Arp2/3 with CK666 did not change the connectivity of the actin cortex for suspended cells, while it did so strongly for the adhered cells (Figure S5B). Therefore, we assume that the role of Arp2/3 as a crosslinker is reduced in suspended cells, which is reinforced by the down-regulation of Rac1 and Cdc42 in suspended cells. This means that the actin cortex of suspended cells consists of short and weakly crosslinked actin filaments compared to that of adhered cells. This might explain why the stiffness of suspended cells is significantly lower than the stiffness of adhered cells even though MHA is decreased. Consequently, the activation of phospho-cofilin via the intrinsic down-regulation of ROCK appears to decouple MHA from the stiffness of suspended cells. Moreover, the tension in the actin cortex of suspended cells has been shown to be significantly lower than that of adhered cells [2], which might also be explained by the absence of (high tension) stress fibers in suspended cells. Both of these explanations support our hypothesis of the intrinsic down-regulation of ROCK in suspended cells.

Nonetheless, there remains an open question: When a decrease in actin filament length is expected upon Y-27632 treatment (see above), why does Y-27632 treatment lead to significantly increased actin filament length for the over the nucleus and the perinuclear regions of adhered cells? The SEM images (Figure S6) revealed that after Y-27632 treatment what remained was mainly actin filament bundles, which are naturally more stable than single actin filaments (i.e., straighter, longer). This is best seen for the perinucleus region in Figure S6. It has been shown that higher tension in actin filaments prevents, or at least delays, their severing by cofilin [16]. Similar mechanosensitive trends have been observed elsewhere [17, 18]. Therefore, we would assume that elevated phospho-cofilin activity leads not only to severing of the actin filaments, but also to increased depolymerization of single actin filaments, as supported by the study of Wioland et al. (2017). As we have seen that treatment of these hTERT-RPE1 cells with high concentrations of Y-27632 (i.e., 100 µM) leads to their detachment (data not shown), we can assume that the Y-27632 concentration used reflects an initial state of the transition from adhered to suspended cells. In the further processes, actin bundles and stress fibers are going to be depolymerized to completely re-arrange the actin cortex in suspended cells.

In summary, the actin cortex intrinsically assumes separate *steady-states* for these NPS states. On the one hand, extrinsic alterations to the actin cortex within the NPS states leads to predictable correlations with the DSM measures (Figure 3). On the other hand, intrinsic alterations to the actin cortex through changes in the cell adhesion state promote fundamentally different characteristics of the actin cortex, which depend on the underlying biological processes of the cells, as explained above. Consequently, the actin cortex is a material that has intrinsically different properties, although each of the NPS states of the actin cortex react almost identically to extrinsic alterations.

**NMII localization**

For visualization of the co-localization of actin with NMII, the *Co-localizationAnalysis* tool from Image J was used (Figure 5C, right-hand images), which indicated the actin and NMII co-localized signals in white.

*Our observations on the DSM measures upon NMII interactions (Figure 2) are in line with earlier observations, such as those of Xia et al. (2015), who showed only small effects of blebbistatin on the structure of the actin network, and indeed, no effects on the cell mechanics over the nucleus (Xia 2015). Similar small effects on cell mechanics have been shown in a number of recent studies [19, 20]. Chugh et al. (2017) also showed that inhibition of myosin using blebbistatin did not influence actin cortex thickness, which would otherwise be a good candidate to explain the changes in cell mechanics [21].*

*Indeed, there are a number of other studies that have provided ambiguous, or even contrary, data after interference with myosin activity. A study by Rigato et al. (2015) showed alterations in the actin cortex and the stiffness of blebbistatin-treated RPE1 cells when measured in the perinuclear region, but not when measured over the nucleus [20]. Adhered cells such as NRK cells [22], 16HBE14o cells [23], and HFFs [24] have shown softening after treatment with blebbistatin. This cell softening was also shown when the viscoelastic properties of suspended 3T3 fibroblasts were investigated via bead attachment [25], but it was lost when the cells were fully in suspension, without any bead attachment, where the cells also showed stiffening under myosin inhibition [26]. Obviously, all of these data have been obtained using different cell types and different methods on different time scales. However, we believe that experimental differences will only partially explain the extremely varied, and even opposing, data in the literature.*

*Our observation that NMII minifilaments are not co-localized with the actin cortex does not mean that there is no NMII at all inside the actin layer of the cortex. It is still possible that monomeric NMII is located within this structure, as the small size of monomeric NMII would allow it to penetrate this dense actin meshwork. Nevertheless, as we did not detect NMII within the actin cortex itself using expansion microscopy, we can deduce that the proportion of monomeric NMII in the actin cortex layer is either very low or below the detection limit here. The NMII that we did detect was located just next to the inner side of the actin cortex. It is highly likely that this myosin is present in the form of MNII minifilaments, for two reasons: due to the high density of myosin heads and the associated stained myosin light chains, we would expect a high fluorescent signal, as was detected; and secondly, these NMII minifilaments are relatively large, with a length of ~250 nm and a diameter of ~100 nm [27-29]. As describe above, we do not expect such large filament bundles to enter the actin cortex, as this has a mesh hole diameter of 55 nm to 75 nm. Therefore, NMII minifilaments would only penetrate a little into the actin cortex. This was confirmed by our measurements of a penetration depth of NMII into the actin layer of about 10%.*

*Interestingly, as well as measurement time-scales and their viscoelastic implications, this organization of NMII beneath the actin cortex might explain some of the discrepancies in the literature concerning the effects of NMII-targeting compounds on the actin cortex. Local measurements that form only a small indent in the cell might only minimally deform the actin cortex. Such measurements would therefore not be sensitive on any NMII-targeting treatments. Our AFM measurements presented here are an example of such measurements. However, if AFM is used to form larger indents in cells, any effects of NMII should become apparent. These effects should start to appear when an indentation greater than the thickness of the cortex, >400 nm, is used (see Figure S7). By using AFM, we would expect such an effect to become apparent from a set-point force of ≥1 nN. However, with such a large indent over the nucleus, the potential deformation of the nucleus will start to have a major role. The same argument applies in the perinuclear region where the influence of the substrate (glass) will not be neglectable for large indentations. The actin and myosin arrangement that we propose here might explain other cell mechanics measurements for which the NMII inhibition has shown particular effects: For example, when using microfluidic deformation devices [26], the whole cell is deformed. In this case, the myosin layer will be affected by the global cell deformation, and so effects of NMII inhibition will be observed.*

*We also saw that the polymerization dynamics of the actin cortex changed upon NMII inhibition. Such an effect has been seen before [16, 17], when it was suggested that the severing activity of cofilin was reduced by cellular tension. As blebbistatin reduces the tension of the actin cortex [24], this effect might result in increased cofilin activity, followed by greater actin severing and faster actin dynamics, which is what we have shown here. This idea was also supported by* in-vitro *data from Sonal et al. (2018), where NMII was shown to promote actin turnover* in vitro *[30].*

**Arp2/3 and Formin**

In adhered cells, the structure of the actin cortex over the nucleus that was assessed using SEM was strongly affected by inhibition of Arp2/3, as the treatment with CK666 resulted in ~50% increase in MHA (**Figure 2A**). However, in the perinuclear region, MHA was slightly reduced with this treatment, albeit not significantly. These changes are in line with the changes in the mechanical properties. Indeed, over the nucleus, the stiffness of the cell cortex decreased with CK666, while there was no significant change in the perinuclear region. Finally, inhibition of the Arp2/3 complex increased the actin half-time recovery over the nucleus by a factor of three. In a similar manner as for the other DSM measures, the actin recovery time in the perinuclear region was not significantly different from the control, even though it was approximately three times higher in the nuclear region. In summary here, Arp2/3 inhibition in adhered cells strongly affected the DSM measures of the actin cortex in the nuclear region, but had no significant impact in the perinuclear region.

In parallel to these effects of Arp2/3 inhibition, inhibition of formins increased MHA and decreased the stiffness of the actin cortex for both the over the nucleus and the perinuclear regions of these adhered cells. The actin half-time recovery was also significantly increased over the nucleus, but only showed a trend towards an increase in the perinuclear region. In summary, formin inhibition in these adhered cells strongly affected the DSM measures of the actin cortex in both the over the nucleus and the perinuclear regions.

Immunofluorescence images of adhered cells showed that Arp3 and formin have high signal intensities that co-localize with the nuclear region (Figure S5C-E). The averaged signals here for Arp3 (for 15 cells) and formin (for 8 cells) showed that their radial intensities decreased gradually towards the edges of the cells (Figure S5F, G). Taken together, these results suggest that formin activity in adhered cells is required for the actin cortex in both the over the nucleus and the perinuclear regions, while Arp2/3 activity is only required close to the nucleus.

*It has been suggested that Arp2 and Arp3 are both found in the nucleus [31], although other studies have suggested that the nucleation of the Arp2/3 complex onto actin filaments is favored along curved membranes (as seen for the over the nucleus region compared to the perinuclear region) [3, 4]. Also, although formin distribution and localization appears similar to that of Arp3 (Figure S5D), we do not observe stronger structural or dynamic changes between the over the nucleus and the perinuclear actin cortex regions. Formins are essential for rapid actin polymerization, and as such, it is not surprising that they have importance in the properties of the actin cortex, independent of the location. A lack of formins results in decreased total filament content in the actin cortex, and thus reduced actin density, which will result in increased MHA, as observed here (Figure 2A). Additionally, a decrease in actin filament density induces a reduction in cell stiffness, which has also been reported previously, along with slowing down of the actin dynamics [32, 33].*

Surprisingly, when the cells were in suspension, inhibition of Arp2/3 and formins did not induce any large changes. From these data here, for CK666-treated cells, only the stiffness was affected, while the dynamics and structural properties of the actin cortex were not (Figure 2). Moreover, there were no significant changes in the actin cortex properties of Smifh2-treated cells (Figure 2). Immunofluorescence images of suspended cells showed co-localization of Arp3 and formin with the cortical actin signal (Figure S5A, B).

*Regulation of the Rho family proteins in cells in suspension remains unclear. It has been shown that Cdc42 and GTP-Rac require integrin signaling to correctly anchor to cell membranes [34, 35]. This anchoring is essential for their activity, and as both proteins are upstream of Arp2/3, we can hypothesize that the activity of Arp2/3 will be altered as soon as cells are taken into suspension. Similarly, it has been shown that the activation of ROCK by Rho is mediated by cell tension [5], and it has also been suggested that activation of mDia1 requires such mechanical signaling [36]. The lack of integrin signaling and tension is then also detrimental to the formin activity. Additionally, in a more recent study [37], it was shown that for cells in suspension, formin-mediated filopodia formation was impaired in the absence of Arp2/3. Taking these studies together, this suggests that when cells are taken into suspension and in the absence of cell tension, the activity of both Arp2/3 and formin are strongly reduced. This would explain why their inhibition has less impact, if any, on the properties of the actin cortex of suspended cells.*

### SI Figures

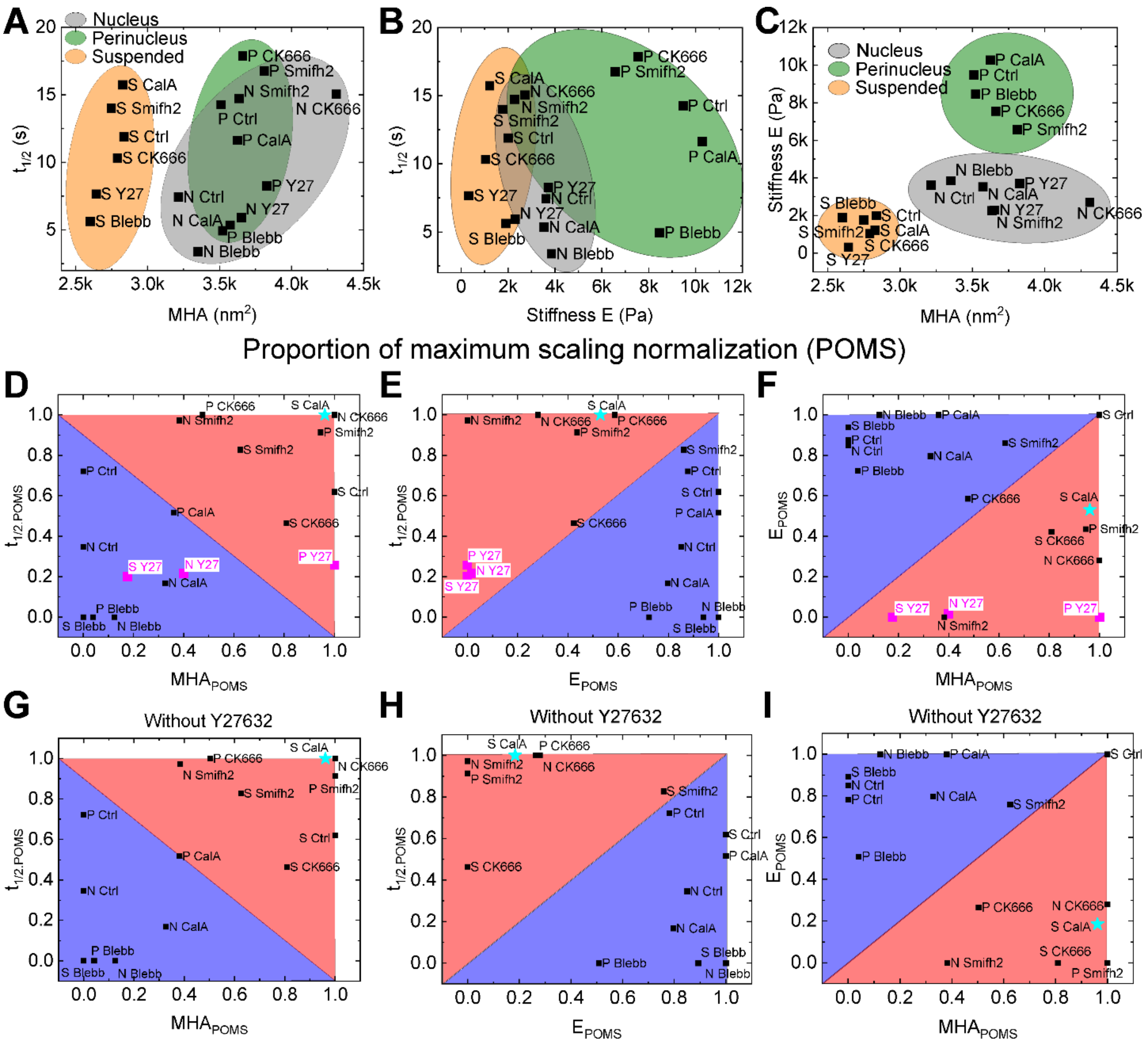


**Figure S1: Distribution of cell states nucleus, perinucleus and suspended cells (NPS) for dynamics, structure and mechanics (DSM) comparisons and POMS normalization.** Distributions of NPS in relation to DSM (**A-C**). Ellipses correspond to one adhesion state: nucleus (gray), perinculeus (green) or suspended cells (beige). Proportion of maximum scaling (POMS) normalization per panel of Figure 2 with Y27632 (**D-F**) and without (**G-I**) Y-27632. Colored triangles represent a guide for the eye to separate between NMII related perturbants (Blebbistatin and CalyculinA) plus Ctrl and actin related perturbants (CK-666 and Smifh2). Y-27632 is marked with white rectangles. Unexpected localizations within diagrams are marked with a light-blue star (CalyculinA in suspended cells only). Color code: Red: Actin related perturbants, blue: NMII related perturbants. For statistics, please see Figure 2. For clarity error bars are not shown, since they are depicted in Figure 2.

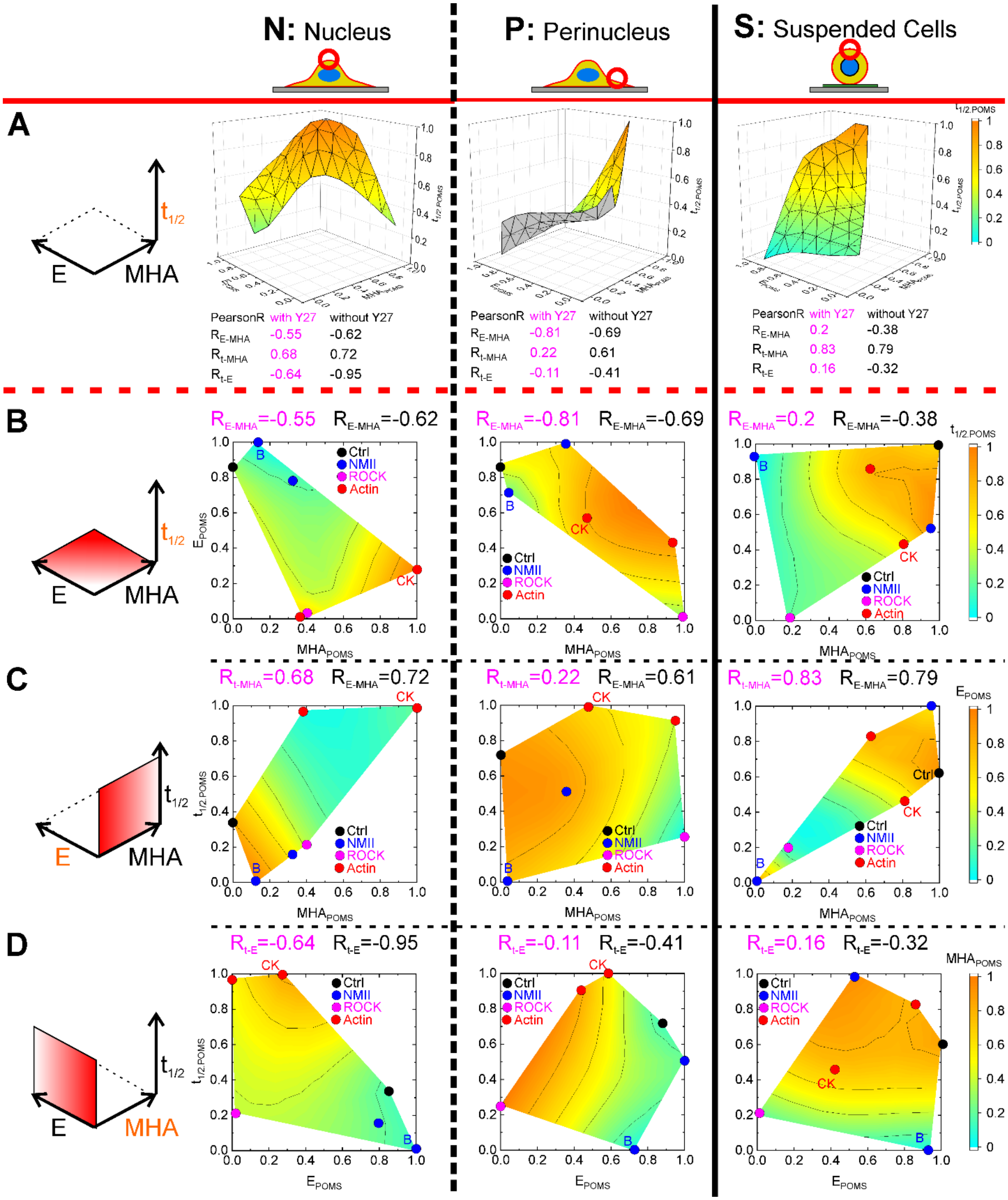


**Figure S2: Correlations between actin dynamics, cortical structure and cellular sechanics upon extrinsic cortex alterations.** 3D correlation diagrams (**A**) are identical to Figure 3 (A-C). For clarity reasons each plane of 3D diagrams is depicted in 2D with third parameter color-coded (from top to bottom for nucleus, perinucleus and suspended cells) (**B-D**). Data points represent the means of each pharmacological perturbant condition ("NMII" refers to Blebbistatin indicated by "B" and CalyculinA, indicated by the color blue without a letter; "ROCK" refers to Y-27632; "Actin" refers to CK-666 indicated by "CK" and Smifh2 indicated by the color red without a letter). Single Pearson correlation coefficients resulted linear approximations of the presented 2D diagrams. Color-codes (and surfaces in 3D and 2D) result from extrapolation between data points.

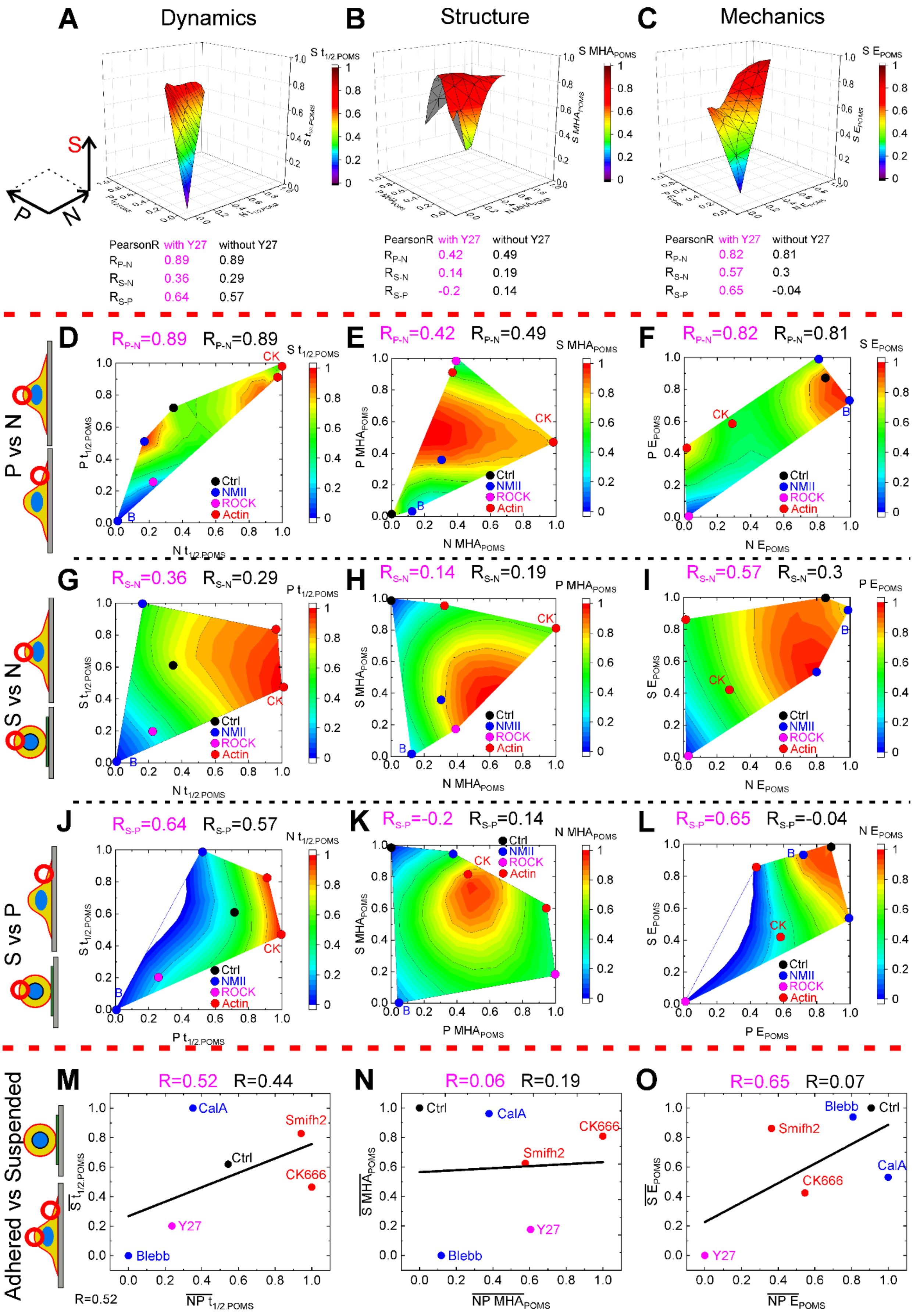
A
Dynamics
B
Structure
C
Mechanics
PearsonR with Y27 without Y27
RP-N 0.89 0.89
RS-N 0.36 0.29
RS-P 0.64 0.57
PearsonR with Y27 without Y27
RP-N 0.42 0.49
RS-N 0.14 0.19
RS-P -0.2 0.14
PearsonR with Y27 without Y27
RP-N 0.82 0.81
RS-N 0.57 0.3
RS-P 0.65 -0.04
P vs N
D RP-N=0.89 RP-N=0.89
E RP-N=0.42 RP-N=0.49
F RP-N=0.82 RP-N=0.81
S vs N
G RS-N=0.36 RS-N=0.29
H RS-N=0.14 RS-N=0.19
I RS-N=0.57 RS-N=0.3
S vs P
J RS-P=0.64 RS-P=0.57
K RS-P=-0.2 RS-P=0.14
L RS-P=0.65 RS-P=-0.04
Ctrl
NMII
ROCK
Actin
CK
B
Adhered vs Suspended
M R=0.52 R=0.44
N R=0.06 R=0.19
O R=0.65 R=0.07
CalA
Smifh2
CK666
Y27
Blebb
R=0.52

**Figure S3: Correlations upon extrinsic cortex alterations comparing nucleus, perinucleus and suspended cells for each parameter of DSM.** 3D correlation diagrams (**A-C**) are identical to Figure 3 (D-F). For clarity reasons each plane of 3D diagrams is depicted in 2D with third parameter color-coded (from top to bottom for dynamics, structure and mechanics) (**D-L**). Data points represent the means of each pharmacological perturbant condition ("NMII" refers to Blebbistatin indicated by "B" and CalyculinA, indicated by the color blue without a letter; "ROCK" refers to Y-27632; "Actin" refers to CK-666 indicated by "CK" and Smifh2 indicated by the color red without a letter). Single Pearson correlation coefficients resulted linear approximations of the presented 2D diagrams. Color-codes (and surfaces in 3D and 2D) result from extrapolation between data points.

Data obtained on the cortex over the nucleus and perinucleus were averaged (as well as re-normalized) and compared to suspended cells for dynamics represented by $t_{1/2}$ (**M**), structure represented by MHA (**N**) and mechanics represented by E (**O**). Pearson R correlations coefficients are calculated with (magenta) and without (black) Y-27632. Data from Figure 2 were normalized using POMS normalizations. For statistics, please see Figure 3 for 3D (A-C) representation and Figure 2 for D-O.

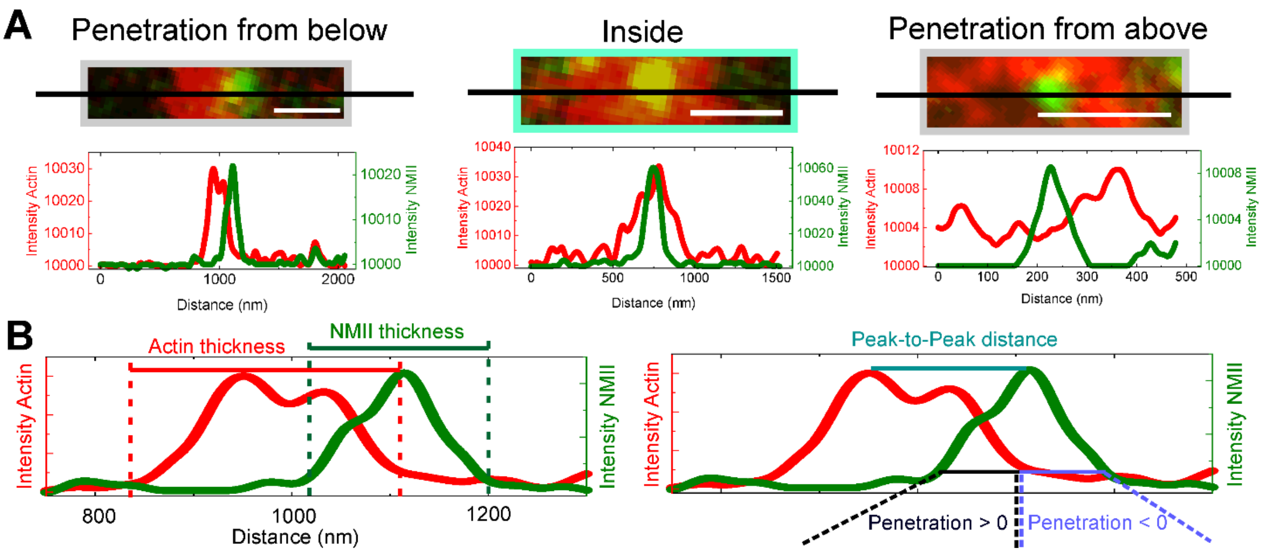


**Figure S4: NMII minifilaments localization analysis.** Identical to Figure 4 (first row) with intensity profiles below (**A**) and parameter illustration on the intensity profile of "Penetration from below" magnified (**B**).

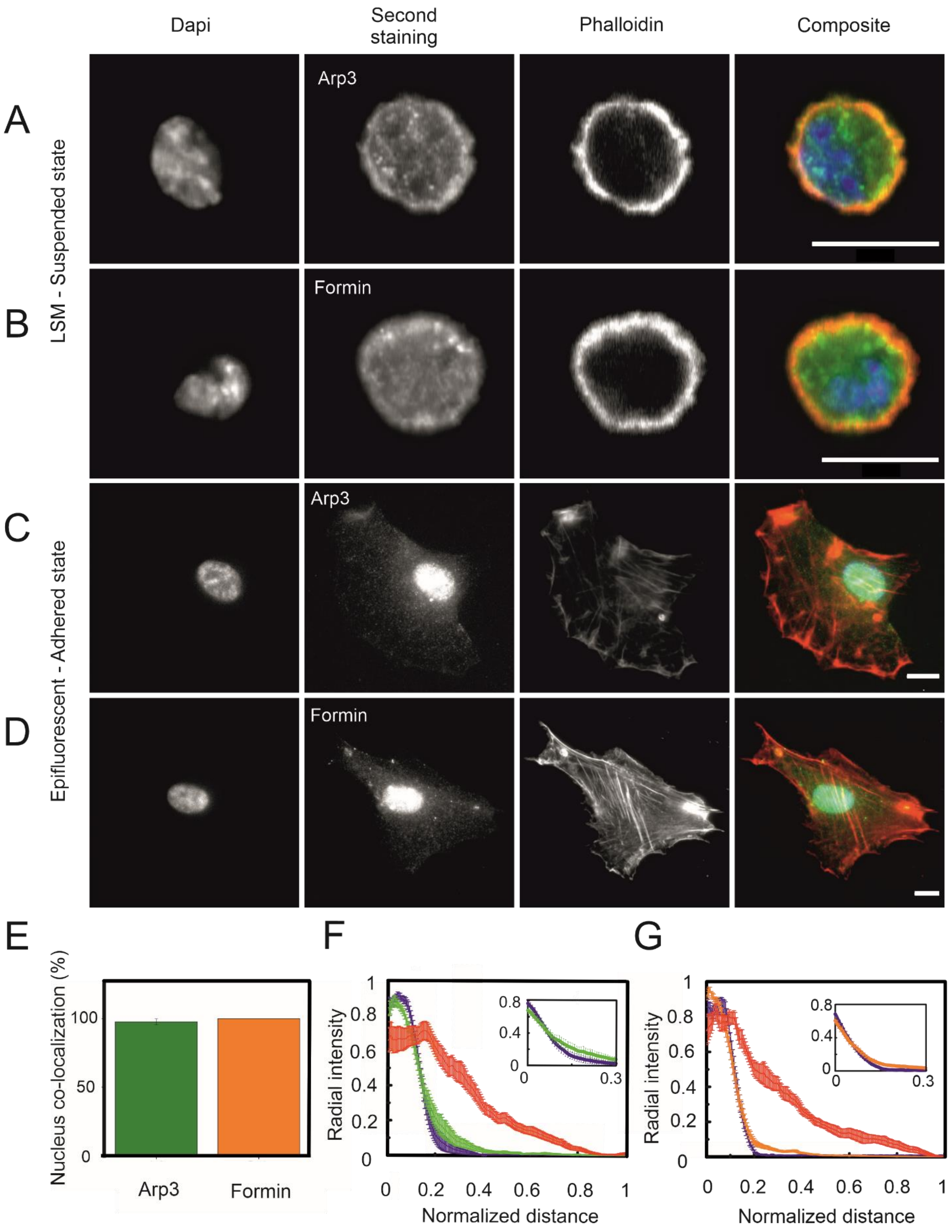


**Figure S5: Localization investigations of Arp3 and Formin.** Arp3 and Formin are depicted in green (second panel) on suspended cells (**A, B**) and adhered cells (**C, D**) as well as nucleus (first panel, blue in composite) and actin (third panel, red in composite). Co-localization of Arp3 and Formin with the nucleus were measured by eye using EPI-fluorescence microscopy (**E**). Representative distributions of Arp3 in green (**F**) and Formin in orange (**G**) within adhered cells were depicted in comparison to actin (red) and nucleus (blue). Scale bars (A-D): 10 µm. Statistics: (E) n= 298 (Arp3), n= 689 (Formin); (F) n= 15; (G) n= 8.

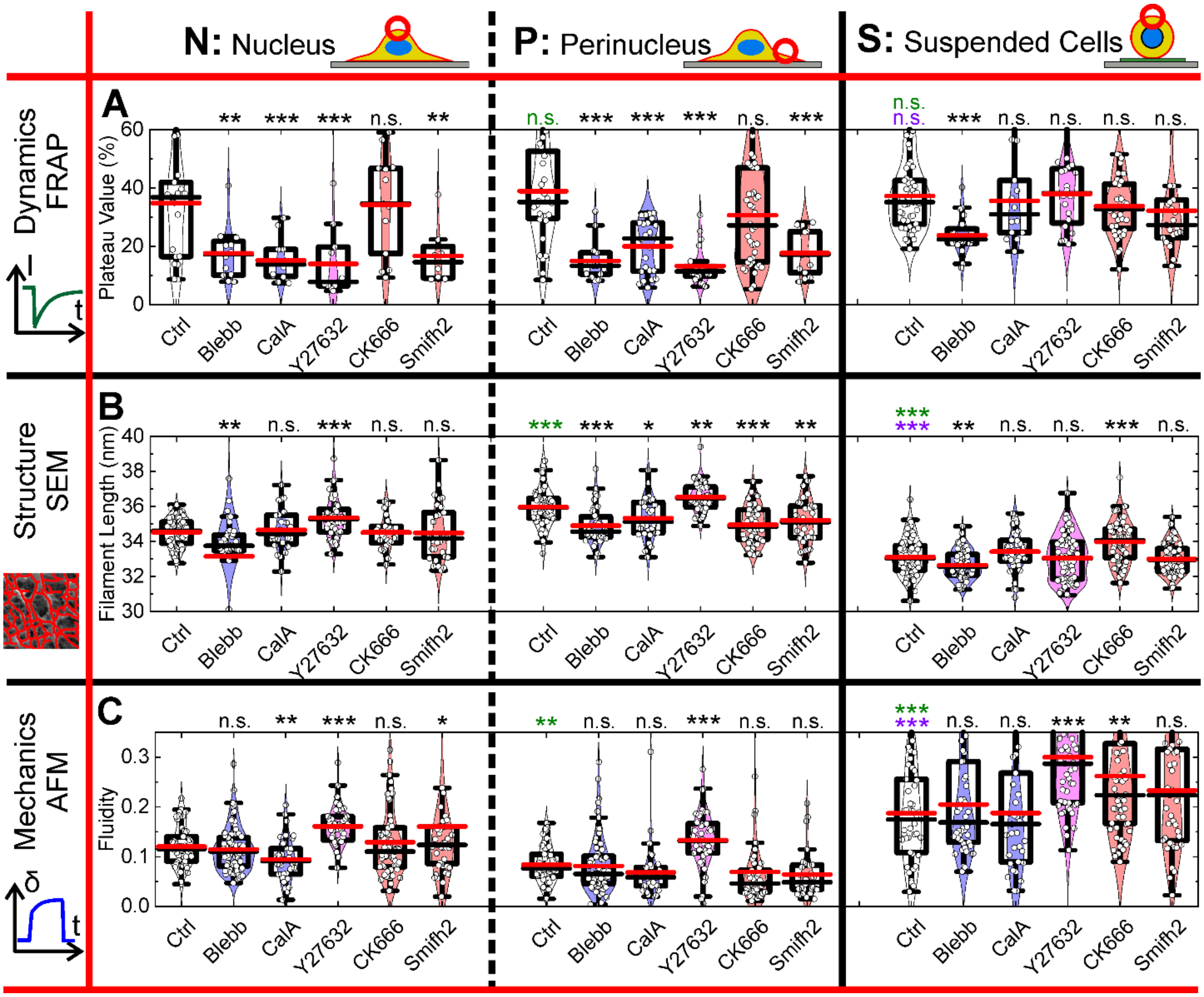


**Figure S6: Structure, mechanics and dynamics of the cellular cortex of adhered (nucleus and perinucleus) and suspended hTERT-RPE1 cells.** Plateau value obtained by FRAP (**A**). Actin filament lengths obtained by SEM was quantitatively analyzed using the algorithm FiNTA by applying a break-off angle of ~17.2° (0.3 rad) (**B**). The fluditiy was quantitatively analyzed employing creep compliance measurements using AFM (**C**). Median values are marked in red, mean values in black. Star method is representing statistical Welch-corrected t-tests. Black stars compare pharmagolocial perturbants with the Ctrl for each panel. Green stars compare controls to Nucleus Ctrl, purple stars compare Suspended Ctrl to Perinucleus Ctrl. n.s.: not significant, *: p<0.05, **: p<0.01, ***: p<0.001.

Cell numbers n are equal to Figure 2 for Dynamics, Structure and Mechanics.

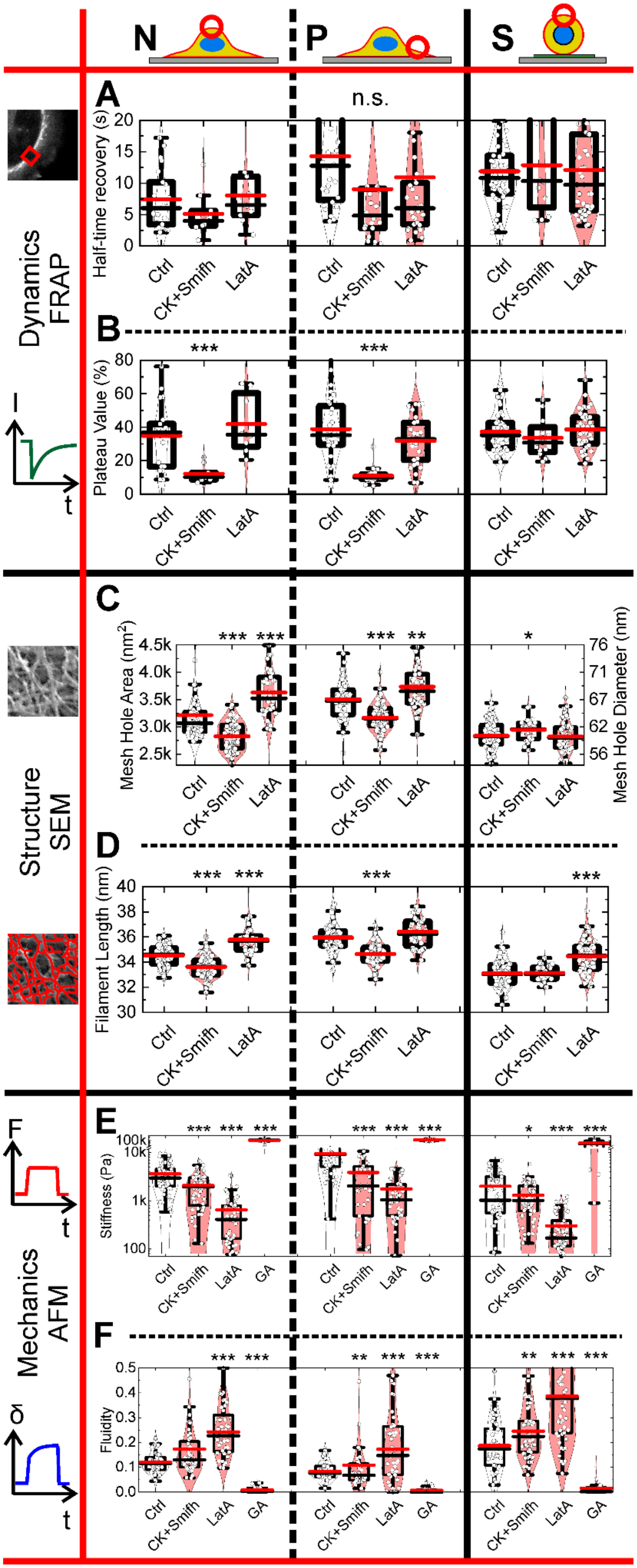

N
P
S
Dynamics FRAP
Structure SEM
Mechanics AFM
A
B
C
D
E
F
n.s.
Half-time recovery (s)
Plateau Value (%)
Mesh Hole Area (nm²)
Mesh Hole Diameter (nm)
Filament Length (nm)
Stiffness (Pa)
Fluidity
Ctrl
CK+Smifh
LatA
GA
I
t
F
σ

**Figure S7: Additional investigations for CK666+Smifh2, LatrunculinA and Glutaraldehyd (AFM only).** Additonal parameters corresponding to parameters of Figure 2 (**A, C, E**) and Figure S6 (**B, D, F**). Median values are marked in red, mean values in black. Statistical tests were performed analogue to Figure 2: Black stars compare pharmagolocial perturbants with the Ctrl of the cell region/state they were investigated in. Green stars compare Controls to Nucleus Ctrl, purple stars compare Suspended Ctrl to Perinucleus Ctrl.

Cell numbers are in the order Ctrl, CK+Smifh, LatA and GA (for AFM only). *Dynamics (FRAP):* Nucleus: n= 8, 9, 8; Perinucleus: n= 13, 22, 29; Suspended cells: n= 23, 12, 41. *Structure (SEM):* Nucleus: n= 31, 88, 37; Perinucleus: n= 33, 82, 44; Suspended cells: n= 70, 27, 61. *Mechanics (AFM):* Nucleus: n= 53, *52, 40, 19; Perinucleus:* n= 52, 34, 43, 17; Suspended cells: n= 42, 50, 54, 15.

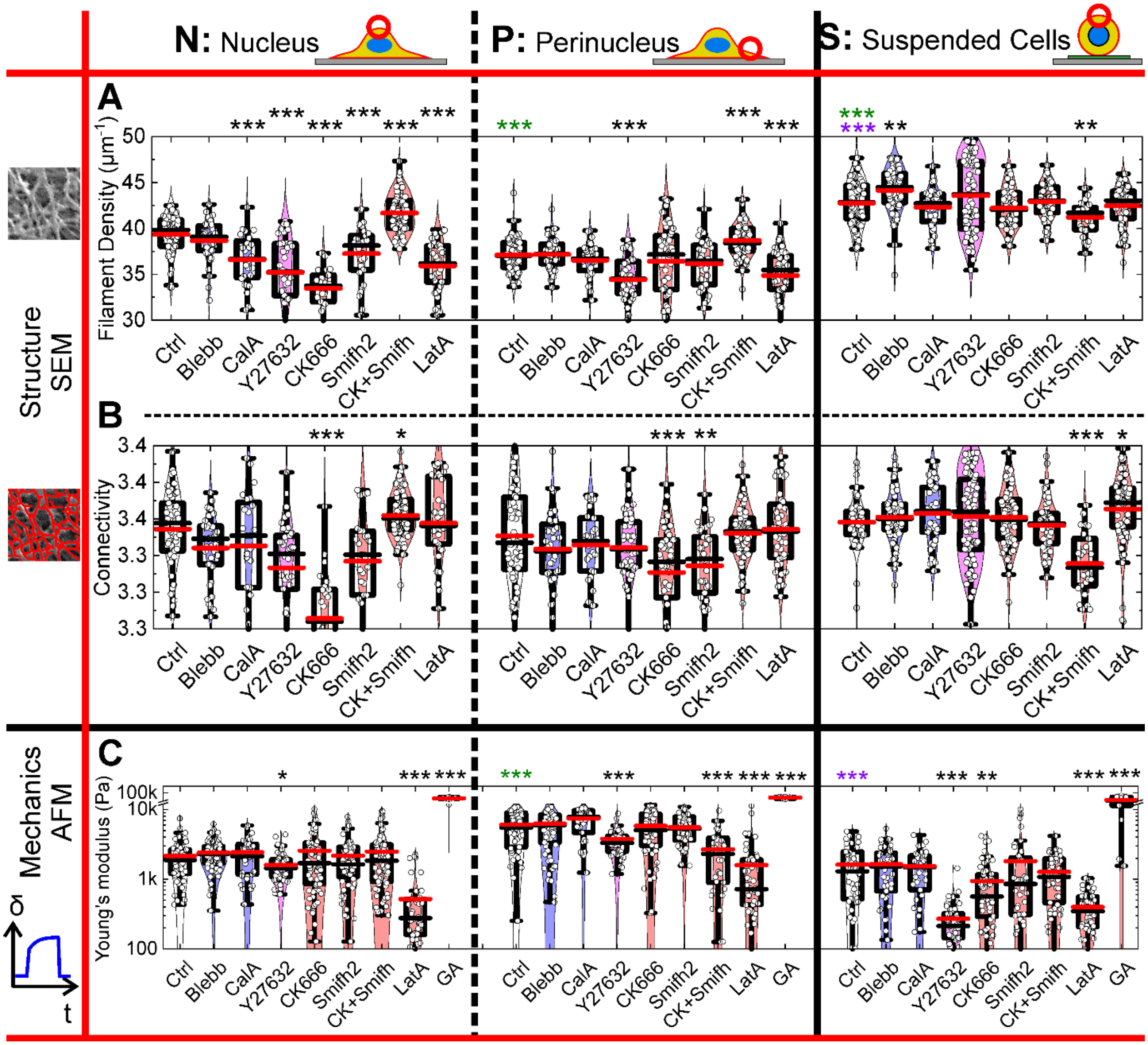


**Figure S8: Additional parameters for structure and mechanics.** Filament density (**A**) and connectivity (so-called coordination number) (**B**) are calculated using FiNTA. Of note, lowest possible value for the connectivity is 3 since 2 would represent a straight line (see **Flormann et al. 2021** for further informations). Young's modulus (**C**) was calculated additionally to stiffness and fluidity. Median values are marked in red, mean values in black. Statistical tests were performed analogue to Figure 2: Black stars compare pharmagolocial perturbants with the Ctrl of the cell region/state they were investigated in. Green stars compare Controls to Nucleus Ctrl, purple stars compare Suspended Ctrl to Perinucleus Ctrl.

Cell number n is equal to Figure 2 for Ctrl-Smifh2 and equal to FigureS3 for CK+Smifh until LatA (or GA, for AFM only).

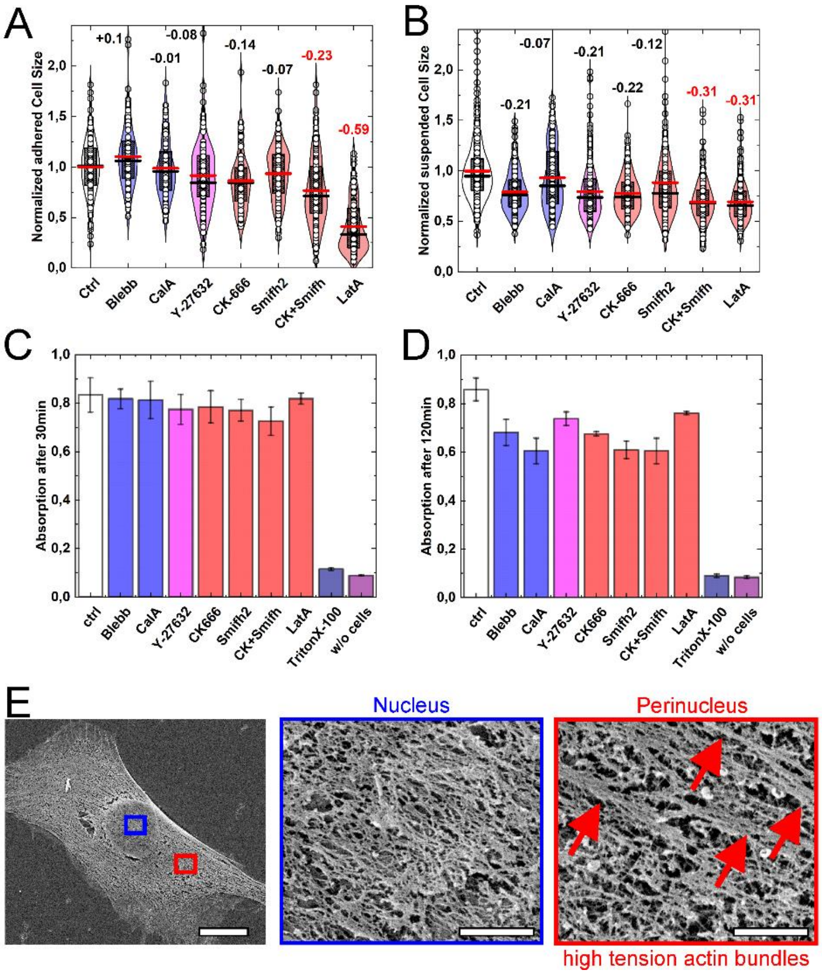


**Figure S9: Cell areas, cell proliferation and visualization of adhered cell regions.** Projected cell areas for adhered (**A**) and suspended (**B**) cells. Median values are marked in red, mean values in black. Mean of the control was normalized to 1. Numbers represent percental deviations from the control mean value for each panel. Red color indicates severe cell shrinkage in comparison to the other deviations within each cellular adhesion state (A, B). No severe cell death upon cytoskeletal perturbators treatment after 30 min (**C**) and 120 min (**D**) was observed using almar blue assays.

Cell numbers n are in the order Ctrl, Blebb, CalA, Y27632, CK666, Smifh2, CK+Smifh, LatA. (A) n= 123, 129, 132, 167, 134, 156, 154, 208. (B) n= 339, 302, 244, 248, 307, 274, 254, 250. (C, D) n= ~30,000 per condition.

SEM image of RPE1 control cell with magnifications of the actin cortex over the nucleus (blue) and at the perinucleus (red) with marked actin bundles (**E**). Scale bars from left to right: 10µm, 1µm and 1µm.

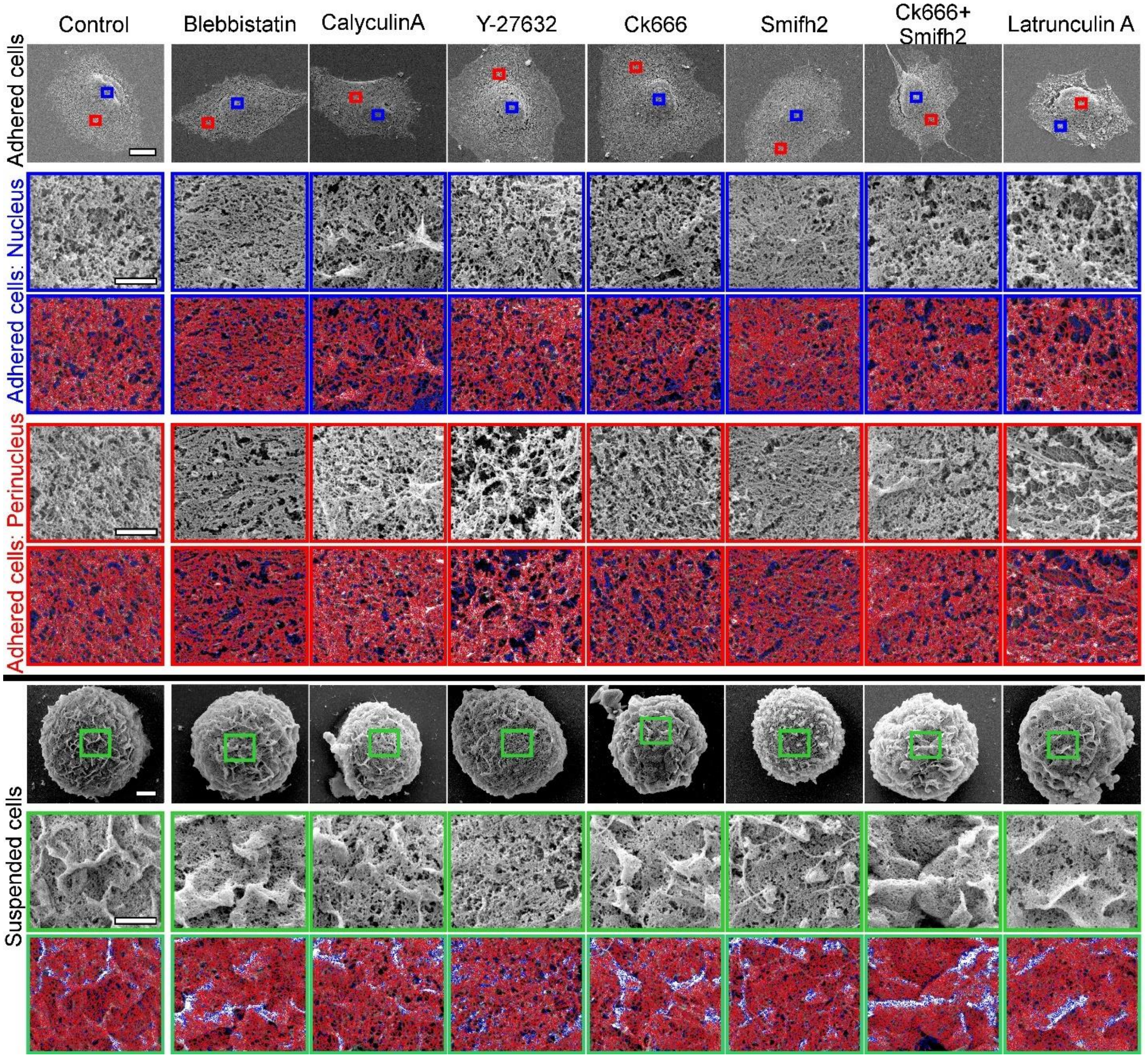


**Figure S10: Representative SEM-images of all pharmacological perturbants in all investigated cell states.** In the first row one cell for each cytoskeletal perturbant is presented. Magnifications of those images are presented below for the cellular cortex over the nucleus (blue) and for the cellular cortex of the perinucleus (red). Analogue representation of suspended cell corticies with magnifications in green. Images were analysed using FiNTA: traced images are directly below untraced images. Tracing lines that were taken into account are depicted in red. Tracing lines that were excluded via intensity thresholding are depicted in blue (intensity thresholding was implemented in FiNTA to reduce tracing error in under- and oversaturated regions of the images). Scale bars from top to bottom: whole cells: 10µm, nucleus: 1µm (identical for traced row), perinucleus: 1µm (identical for traced row), suspended: 2µm, 1µm (identical for traced row). Scale bars are valid for each row.

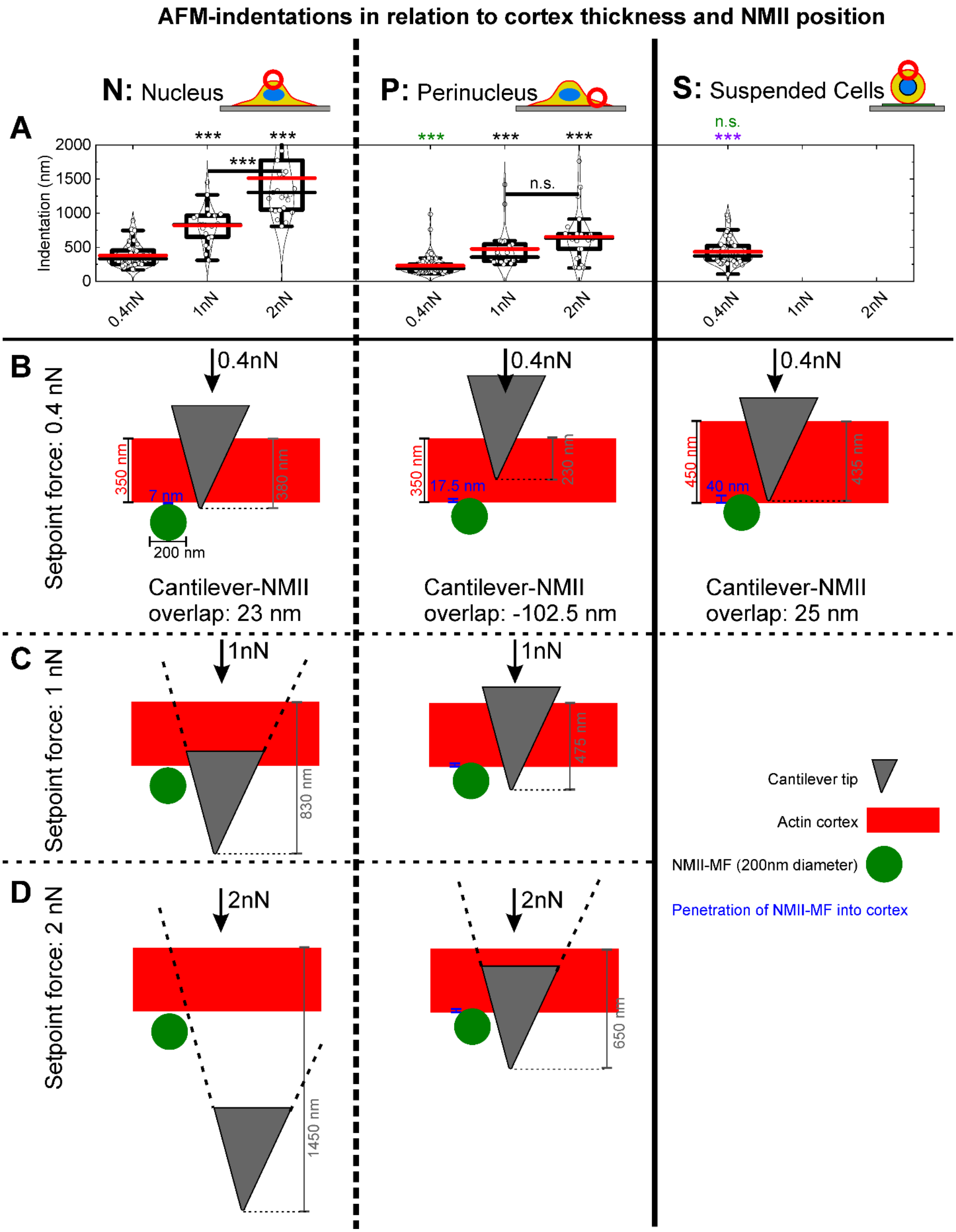


**Figure S11: AFM indentation setpoint force dependency and NMII/actin-localization schemes.** Setpoint force variation on control cells for NPS. Median values are marked in red, mean values in black. Setpoint force of 0.4 nN was used in this study (B). Higher setpoint forces 1nN (C) and 2nN (D) are resulting in larger indentations with maximal indentation for 2nN over the nucleus (D, panel N). Scaling represents measurements of Figure 5 (F, G, I). Suspended cells were not investigated for higher setpoint forces than 0.4nN since high setpoint forces frequently to severe passive cell movements, which was rarely the case for 0.4nN. Potential cortex bending was not visualized since it was not measured in this study explicitly. NMII-MF stands for Non-muscle Myosin II-minifilaments. Black stars compare different setpoint forces to Controls within each state NPS. Green stars compare Controls to Nucleus Ctrl, purple stars compare Suspended Ctrl to Perinucleus Ctrl. n.s.: not significant, *: $p<0.05$, **: $p<0.01$, ***: $p<0.001$.

Cell numbers are in the order 0.4nN, 1nN, 2nN: Nucleus: n= 56, 21, 27; Perinucleus: n= 65, 21, 26; Suspended cells: n= 53.
